# Ketamine’s pharmacogenomic network in human brain contains sub-networks associated with glutamate neurotransmission and with neuroplasticity

**DOI:** 10.1101/2020.04.28.053587

**Authors:** Gerald A. Higgins, Samuel A. Handelman, Ari Allyn-Feuer, Alex S. Ade, James S. Burns, Gilbert S. Omenn, Brian D. Athey

**Affiliations:** Department of Computational Medicine and Bioinformatics, University of Michigan Medical School, Ann Arbor, MI 48109, USA; Phenomics Health Inc., Ann Arbor, MI 48109, USA; Department of Internal Medicine, University of Michigan Medical School, Ann Arbor, MI 48109, USA; School of Public Health, University of Michigan, Ann Arbor, MI 48109, USA; Department of Human Genetics, University of Michigan Medical School, Ann Arbor, MI 48109, USA; Department of Psychiatry, University of Michigan Medical School, Ann Arbor, MI 48109, USA

**Author notes:** Corresponding author: (B.D.A.). GlaxoSmithKline, King of Prussia, PA 19406, USA.

## Abstract

The pharmacogenomic network responsible for the rapid antidepressant action of ketamine and concomitant adverse events in patients has been poorly defined. Integrative, multi-scale biological data analytics helps explain ketamine’s action. Using a validated computational pipeline, candidate ketamine-response genes and regulatory RNAs from published literature, binding affinity studies, and single nucleotide polymorphisms (SNPs) from genomewide association studies (GWAS), we identified 108 SNPs associated with 110 genes and regulatory RNAs. All of these SNPs are classified as enhancers, and additional chromatin interaction mapping in human neural cell lines and tissue shows enhancer-promoter interactions involving other network members. Pathway analysis and gene set optimization identified three composite sub-networks within the broader ketamine pharmacogenomic network. Expression patterns of ketamine network genes within the postmortem human brain are concordant with ketamine neurocircuitry based on the results of 24 published functional neuroimaging studies. The ketamine pharmacogenomic network is enriched in forebrain regions known to be rapidly activated by ketamine, including cingulate cortex and frontal cortex, and is significantly regulated by ketamine (*p*=6.26E-33; Fisher’s exact test). The ketamine pharmacogenomic network can be partitioned into distinct enhancer sub-networks associated with: (1) glutamate neurotransmission, chromatin remodeling, smoking behavior, schizophrenia, pain, nausea, vomiting, and post-operative delirium; (2) neuroplasticity, depression, and alcohol consumption; and (3) pharmacokinetics. The component sub-networks explain the diverse action mechanisms of ketamine and its analogs. These results may be useful for optimizing pharmacotherapy in patients diagnosed with depression, pain or related stress disorders.

**One Sentence Summary:** The ketamine network in the human brain consists of sub-networks associated with glutamate neurotransmission, neuroplasticity, and pharmacokinetics.

## Introduction

Existing antidepressant medications are not effective for many patients (*1*). Novel antidepressant drugs include compounds that target the N-methyl-d-aspartate receptor (NMDAR), glycine receptor, or the α-amino-3-hydroxy-5-methyl-4-isoxazolepropionic acid receptor (AMPAR) in the human brain. For example, ketamine (*RS*-2-chlorophenyl-2-methylamino-cyclohexanone), a noncompetitive NMDAR antagonist first approved by the U.S. Food & Drug Administration (FDA) as a surgical anesthetic in 1970 (*2*), is being increasingly used to treat refractory depression (*3, 4*). There is evidence that the widespread action of ketamine in the human brain impacts a number of pharmacodynamic targets including AMPARs, cholinergic receptors, calcium channels and other neurotransmitter molecules, although most evidence suggests that ketamine and its analogs exert their actions through partial antagonism of the NMDAR in the human brain (*5-8*). One presumptive target for ketamine-like drugs is the NMDAR, which also contains binding sites for glycine and D-serine encoded by *GLRB*, and sites that bind polyamines, histamine, and cations. Neuroimaging studies have demonstrated that intravenous infusion of ketamine causes a transient surge in glutamate levels detected in prefrontal cortex and cingulate cortex in parallel with its rapid antidepressant action (*5, 7*).

A 2018 review provided detailed evidence of the therapeutic mechanisms of ketamine, its enantiomers, and active metabolites (*8*). Research suggests that ketamine’s antidepressant action might result from stimulation of dendritic spine plasticity in the prefrontal cortex (*9*) or through activation of the inflammasome (*10*). Ketamine and its metabolites also act as modulators of the opiate, cannabinoid, and related receptors (*8*), the hyperpolarization activated cyclic nucleotide gated potassium channel (*11*), the estrogen receptor, and the AMPA receptor subunits GRIA1 and GRIA4 (*12*), and many other known (*8, 13*) and unknown pharmacodynamic targets within human brain. Ketamine and its metabolites strongly induce the expression of the *CYP2B6* gene in human brain, which encodes a drug metabolizing enzyme that contributes to first- and second-pass metabolism of the drug and its metabolites (*14*). Recent genomewide association studies (GWAS) in humans demonstrate association of ketamine response and adverse events with enhancers of genes and long non-coding RNAs (lncRNAs) related to the roundabout guidance receptor 2 (*ROBO2*) gene, whose product binds members of the slit guidance ligand family (*SLIT1, SLIT2*) that are involved in dendrite guidance and synaptic plasticity (*15, 16*). Like phencyclidine, a structurally related compound, ketamine induces acute dissociation, with both drugs exhibiting species-specific differences in response. In sum, the central nervous system (CNS) pathway(s) responsible for the rapid antidepressant effects of ketamine and its enantiomers in patients diagnosed with treatment-resistant depression (TRD) remain poorly defined, including emergence of on- and off-target effects and individual differences in response and adverse events.

Problematic adverse events that characterize response to ketamine in humans include acute psychological effects such as dissociation, delirium, and cognitive impairment. The drug’s dissociative psychotropic effects have emerged as a mechanism to explain the analgesic, anesthetic and sedative effects of ketamine (*1*). Both the racemic and enantiomeric formulations of ketamine produce potent and undesirable psychotomimetic adverse events, including acute sensory distortions, derealization, depersonalization, identity confusion, identity alteration, amnesia, hallucinations, anxiety, and fear (*2, 5, 7, 8*). Dysphoric emergence phenomena occur in a dose-dependent manner, while transient phenomena include elevated blood pressure and heart rate, nausea, and vomiting (*6, 8*).

Intravenous and oral formulations of ketamine have demonstrated efficacy and tolerability in controlled trials and in open-label studies across patient populations known to have little to no response from traditional antidepressants (*6-8*). The *S*-enantiomer of ketamine (Esketamine; Spravato™) has been approved by the U.S. FDA as a nasal spray to be used in combination with a serotonin reuptake inhibitor-based antidepressant for patients who have been diagnosed with TRD (*4*). Esketamine has been approved only for administration by a healthcare professional in a clinical setting because it may produce acute adverse events. In sum, ketamine and its analogs are promising therapeutics for patients diagnosed with depression, but it is uncertain how this powerful psychotropic exerts its widespread response and adverse effects in the human brain (*9, 10, 17-21*).

Our strategy analyzed disparate data from multiple scales of biological function combining bioinformatics, computational medicine and machine learning (*22-25*). GWAS SNPs provided insight into the phenotypic consequences of the mutational alterations and variation within disease sub-networks impacted by ketamine, spatial interactions within chromatin that control expression of ketamine-response genes, and mesoscale functional neurocircuits within human brain regions that are rapidly activated following ketamine administration **(Figure 1)**. We hypothesized that the mechanisms of the different on- and off-target effects of ketamine enantiomers could be determined from studies of disease risk and pharmacogenomic variation in the regulatory, non-coding genome or “regulome” (*23, 26*). Most GWAS SNPs are located within enhancers in the three-dimensional (3D) spatial genome and allele-specific open chromatin provides the foundation on which a SNP may impact gene expression and pharmacogenomic variation among humans (*27-29*). This general feature provides a basis for the machine learning algorithms used in this study to determine the putative causal nature of SNPs. If a variant SNP is silenced by heterochromatin or is located within euchromatin on an allele shrouded by chromatin, it cannot permit the activation of gene expression by transcription factors. Recent studies demonstrate that the topology of the human genome determines regions of activation and repression of gene transcription and translation, organized into enhancer-promoter and promoter-promoter loops in chromatin and long range interactions mediated by superenhancers, topologically associating domains (TADs), and A and B compartments in chromatin (*30, 31*).

**Figure 1.**
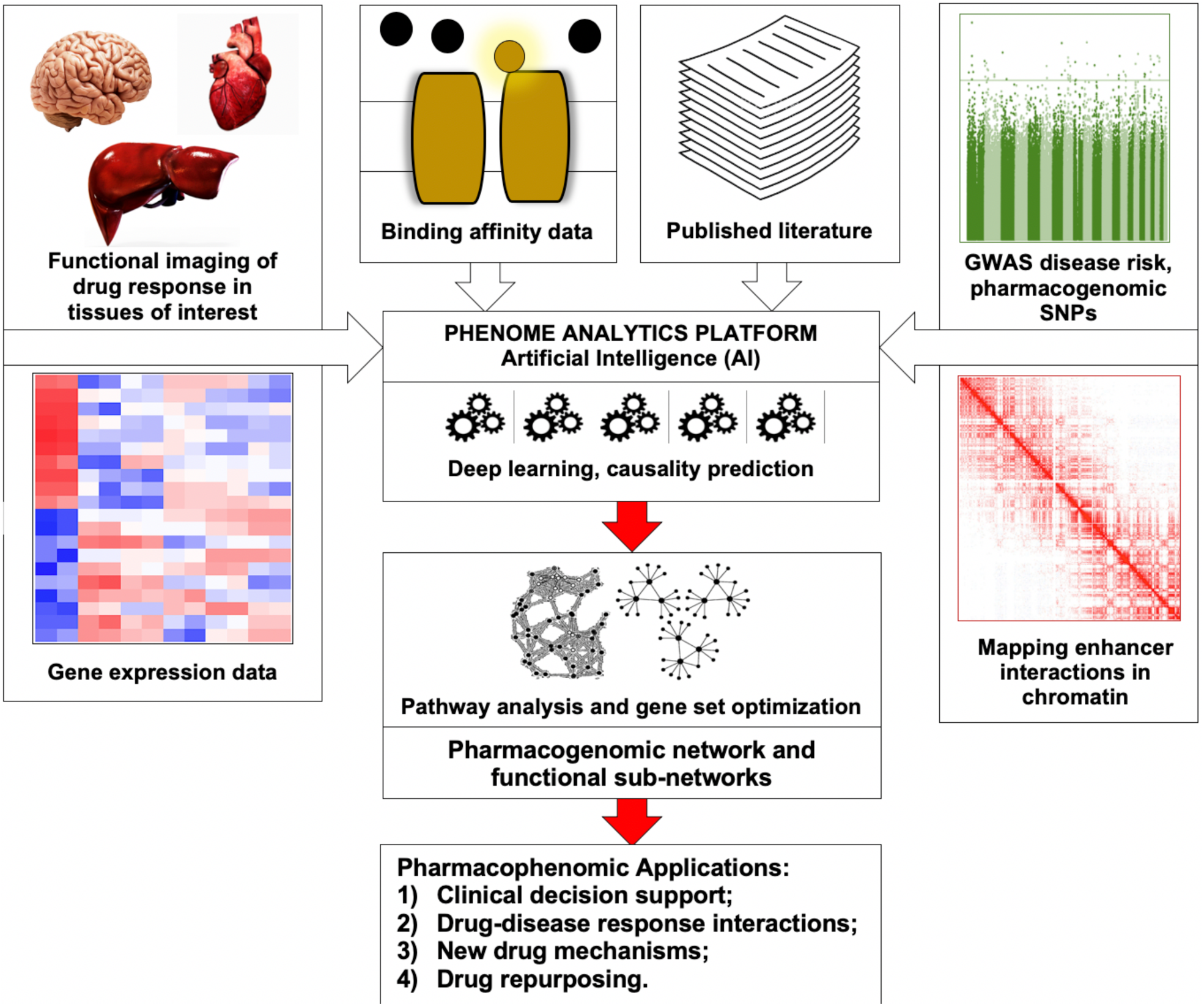
Multi-scale biological data analysis. This strategy for mapping drug networks provides insight into the *mechanistic* on- and off-target effects, laying a foundation for subsequent preclinical studies.

To understand if enhancer networks are active in the same human brain regions where ketamine first exerts a rapid antidepressant response, we determined whether the expressed target genes and their enhancer networks are concentrated within these regions of the CNS. This was accomplished by matching ketamine pharmacogenomic gene expression with higher-order structures where the drug first acts in the human brain, determining whether the GWAS SNPs associated with ketamine’s sub-networks act in these CNS regions, and assessing whether enhancer and superenhancer regulatory elements are localized to this circumscribed neuroanatomical substrate **(Figure 1)**.

## Results

### The ketamine pharmacogenomic network in the human brain

The ketamine pharmacogenomic network we identified consists of 110 genes and regulatory RNAs and exhibits significant overlap with the “cardiovascular disease, neurological disease and organismal injury abnormalities” disease network category as determined using IPA™ (*32*) (*p*=1E-54; Fisher’s exact test). The proteins encoded by the genes in the network exhibit statistically significant STRING sub-network associations (*data not shown*) (*33*). The top 10 most significant biological processes determined by the Panther/Gene Ontology database (*34*) include trans-synaptic signaling, regulation of membrane potential, behavior, response to drug, and nervous system development. The most significant upstream xenobiotic regulator of the network is ketamine at *p*=6.26E-33 (*32, 35*). The top 10 disease gene risk variant categories enriched in the ketamine pharmacogenomic network include cognitive impairment, dissociative disorder, and mood disorders. These results point to a ketamine pharmacogenomic network in the human CNS, because the most significant biological functions of the network are consistent with what is known about ketamine’s mechanisms, including NMDAR and glutamate neurotransmission (*5-9*). The ketamine pharmacogenomic network genes and regulatory RNAs exhibit circumscribed localization within 41 TADs, or less than 2% of human TADs (*36*).

### Component sub-networks of the ketamine pharmacogenomic network

Gene set optimization resulted in 2 distinctly different sub-networks. The glutamate receptor sub-network is enriched for synaptic signaling, glutamate receptor signaling, glutamate pathway regulation and chromatin organization. The top xenobiotic (chemical-drug) up-regulator of the glutamate receptor sub-network is ketamine at p=2.1E-09 **(Figure 2A)** (*32, 35*). In contrast, the neuroplasticity sub-network is enriched for regulation of nervous system development, regulation of neurogenesis, regulation of neuronal differentiation, neurogenesis and nervous system development **(Figure 2B)** (*34*). The neuroplasticity sub-network exhibits significant overlap with the “cardiovascular disease, neurological disease and organismal injury abnormalities” network category as determined by IPA™ (*32*) at *p*=1E-59 and its top xenobiotic up-regulator is also ketamine at *p*=6E-12 **(Figure 4)** (*32, 35*).

**Figure 2.**
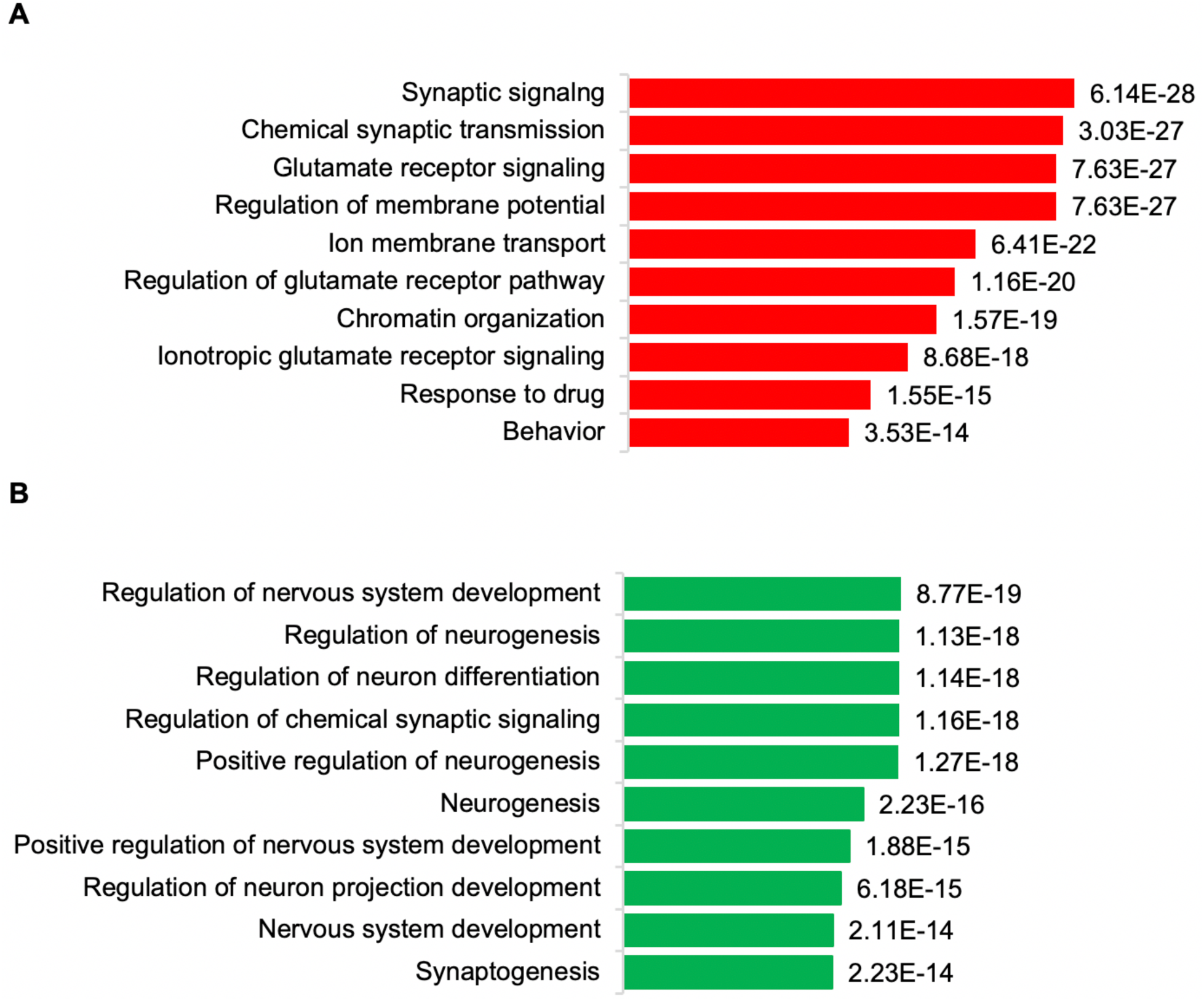
The 10 most significant biological processes from Gene Ontology for the two out of three ketamine pharmacogenomic sub-networks in human brain. (A) The ketamine glutamate receptor sub-network; (B) The ketamine neuroplasticity receptor sub-network. Statistical enrichment using GO (*34*) for “*Homo sapiens*” and “nervous system.”

**Figure 3.**
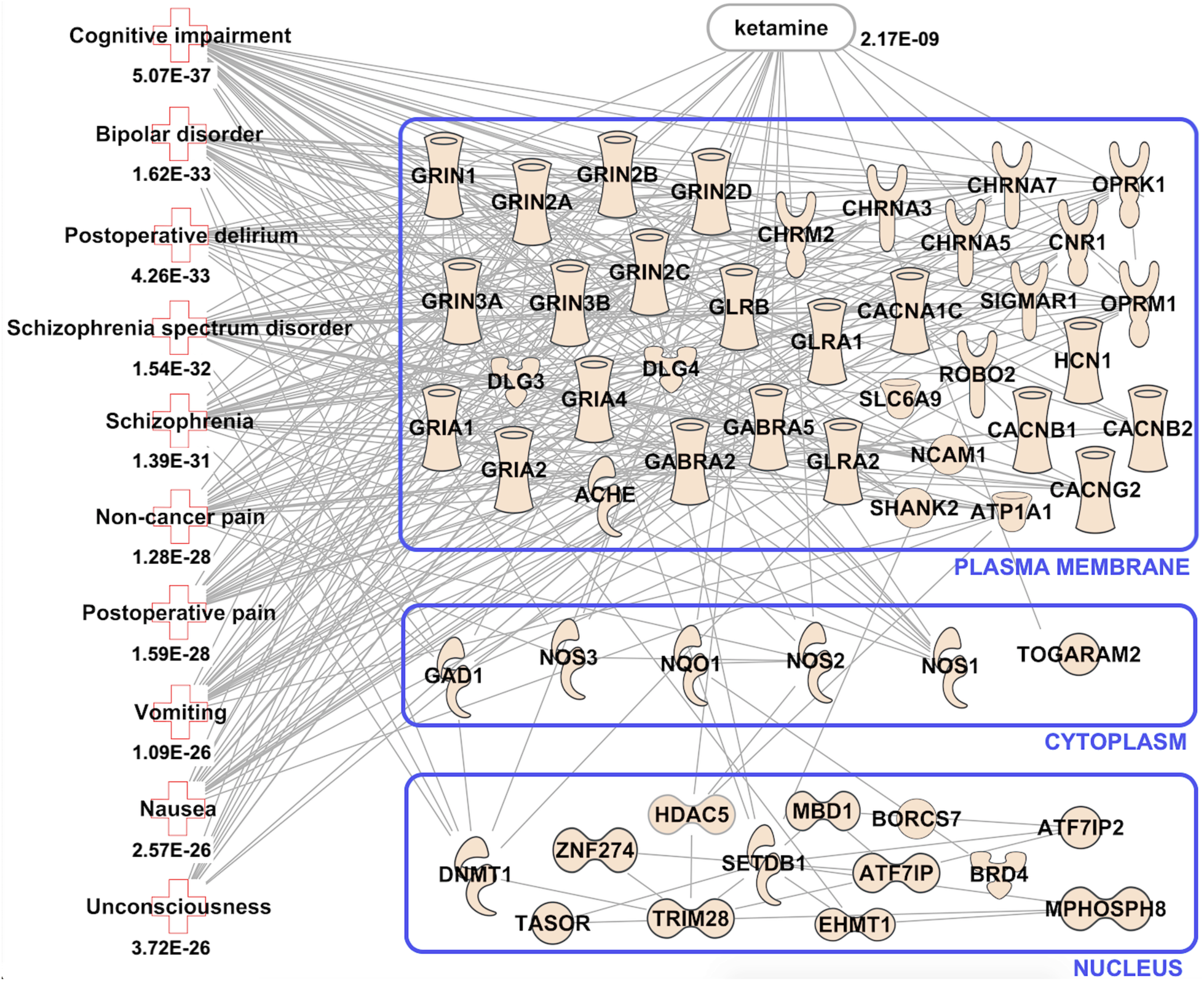
Diseases and conditions associated with the ketamine pharmacogenomic glutamate receptor sub-network, based on the published literature, and visualized using IPA™ (*32*). Lines between the symbols indicate significant interconnectivity as defined by IPA™ (*32*), STRING (*33*) and KEGG (*35*), but do not provide information about the nature of the relationships between genes, their protein products, or regulatory RNAs. **Supplementary Table 2A** lists the gene definitions for the members of the ketamine glutamate receptor sub-network. **Key to gene symbols:**

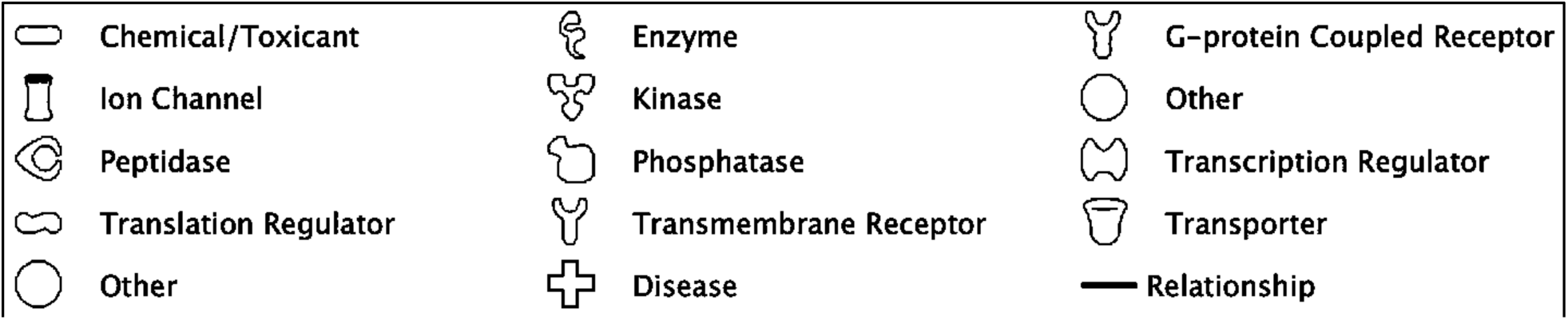

**Figure 4.**
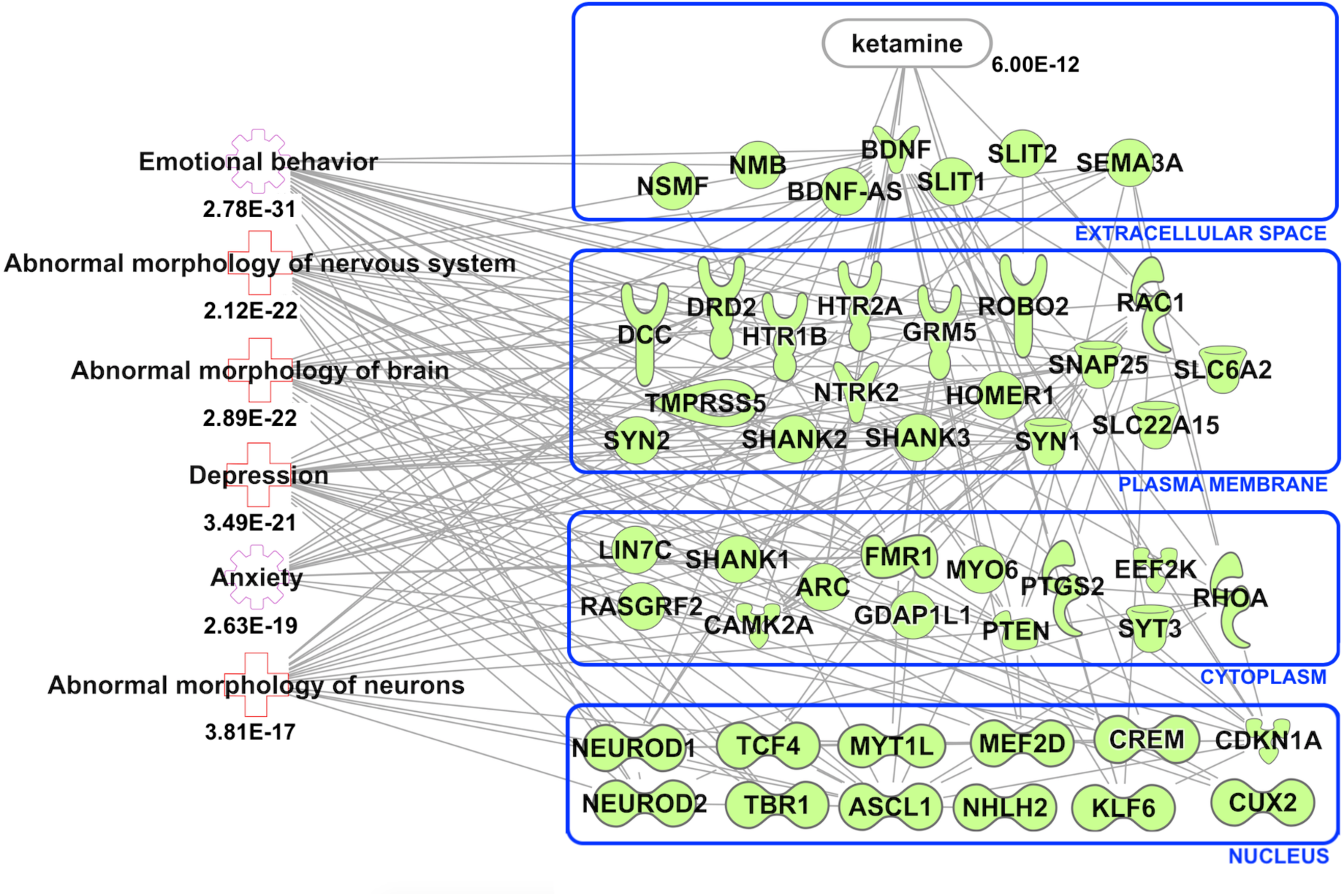
Diseases and conditions associated with the ketamine neuroplasticity sub-network, based on the published literature, and visualized using IPA™ (*32*). Lines between the symbols indicate significant interconnectivity as defined by IPA™ (*32*), STRING (*33*) and KEGG (*35*), but do not provide information about the nature of the relationships between genes, their protein products, or regulatory RNAs. **Supplementary Table 2B** lists the gene definitions for the members of the ketamine glutamate receptor sub-network. The Key for the symbols in this figure is shown in the caption of **Figure 3**.

### The ketamine glutamate receptor sub-network

Our computational pipeline, based on ketamine-response genes selected from published results, showed differences in disease risk annotation between the glutamate receptor sub-network and the neuroplasticity sub-network, even though the products of both sub-networks are regulated by ketamine. Pathway analysis shows that the glutamate receptor sub-network is significantly associated with cognitive impairment, bipolar disorder, postoperative delirium, schizophrenia affective disorder, schizophrenia, non-cancer pain, postoperative pain, vomiting, nausea, and unconsciousness **(Figure 3)**.

Analysis of glutamate receptor sub-network genes using Gene Ontology (*31*) for human CNS biological processes shows the genes *CACNA1C, CACNB2, DLG4, GRIN1, GRIN2A, GRIN2B, GRIN2C, GRIN2D, GRIN3A* are significantly enriched for NMDA glutamate neurotransmission. The genes *SETDB1, TRIM28* and *ZNF274* are enriched for chromatin remodeling in neuronal differentiation, and *BRD1, EHMT1* and *MBD1* are enriched for histone 3 lysine 9 (H3K9) methylation. Different bioinformatic applications (*32-35*) and GWAS disease risk and pharmacogenomic SNP annotations (*37*) showed that essential genes associated with cognitive impairment include *ATFIP, BORCST, GRIA4* and *GAD1*. Mutations in the calcium channel gene *CACNA1C* has been significantly associated with schizophrenia spectrum disorders, including bipolar I disorder, as have *CACNB2, GRIN2A* and *HCN1* **(Supplementary Table 4)**. Post-operative delirium has been associated with the expression of *ACHE, CHRM2, GABRA2, GABRA5, GLRA1, GLRB, GRIA1*, including genes that encode the protein constituents of the NMDA receptor. Nausea and vomiting, common side effects of ketamine therapy, have been associated with cholinergic and opiate gene expression.

### The ketamine neuroplasticity sub-network

The neuroplasticity sub-network is significantly associated with emotional behavior, abnormal morphology of the nervous system, abnormal morphology of brain, depression, anxiety, and abnormal morphology of neurons **(Figure 4)**. Enrichment of sub-network members using Gene Ontology (*31*) for human CNS biological processes associates the genes *ARC, CAMK2A, CNR1, ROBO2, SEMA3A, SLT1* and *SLIT* with synaptic and dendritic plasticity, and *ASCL1, CUX2, BDNF, DCC, KLF6, NEUROD1, NEUROD2, NHLH2, SYN1, SYN2, TBR1* and *TCF4* with neurogenesis.

### The ketamine pharmacokinetic sub-network

A third pharmacokinetic sub-network contains genes that encode cytochrome P450 enzymes, *CYP2B6, CYP2A6* and *CYP3A4*, whose products are responsible for the metabolism of ketamine enantiomers (*8, 12*). This sub-network also contains the *ESR1* gene, which encodes the estrogen receptor alpha subunit (*37*), and *TCERG1*, which encodes a nuclear protein that regulates transcriptional elongation and pre-mRNA splicing (*38*) **(Figure 5)**. The ketamine pharmacokinetic sub-network also contains genes from the other sub-networks, including *ANAPC2, DLG4, EFF2K, GRIA1, GRIA4, GRIN1, GRIN2B, MYO6* and *ROBO2*, so is not significantly different from the other two ketamine sub-networks.

**Figure 5.**
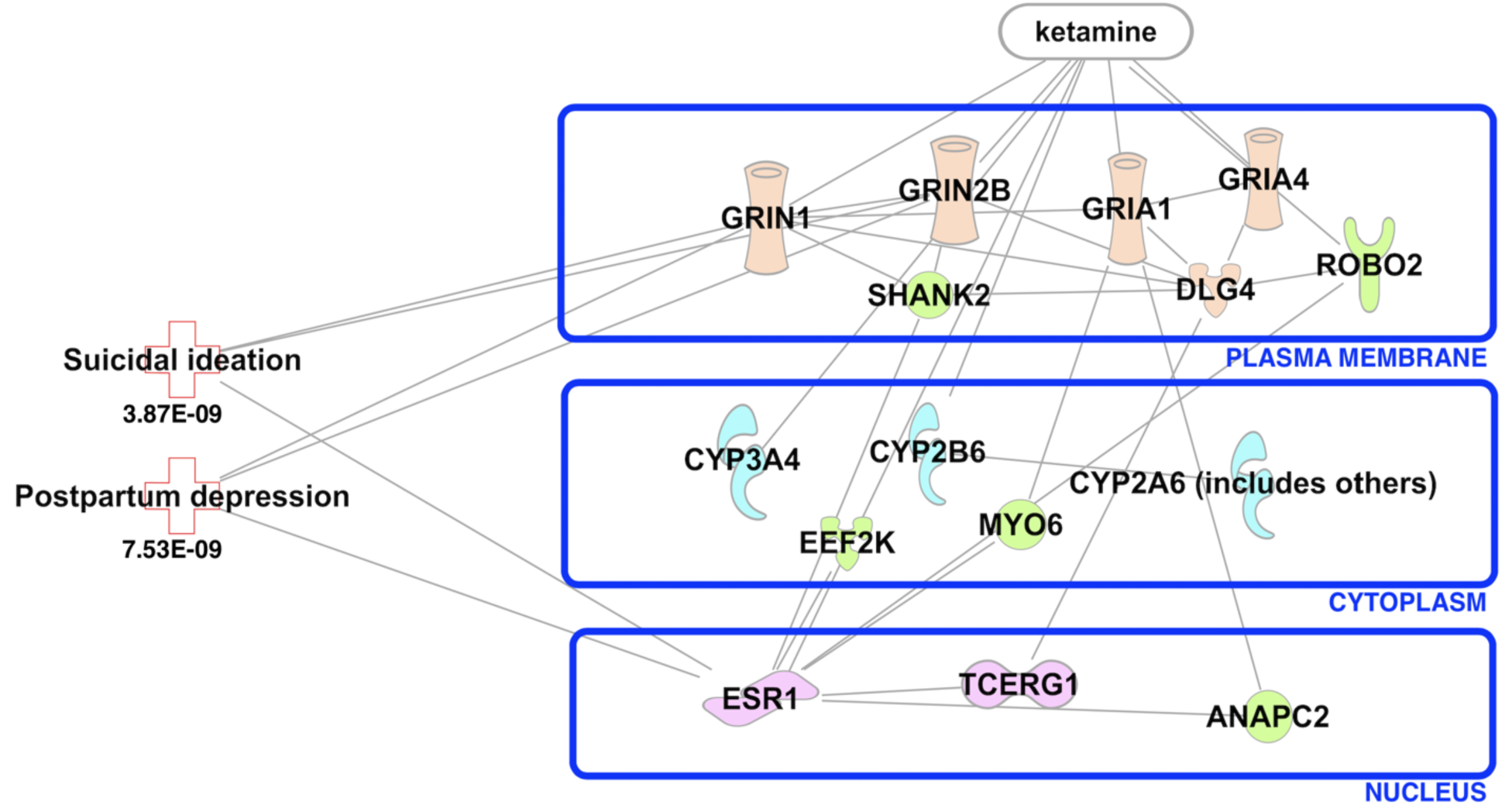
Diseases and conditions associated with the ketamine neuroplasticity sub-network, based on the published literature, and visualized using IPA™ (*32*). Lines between the symbols indicate significant interconnectivity as defined by IPA™ (*32*), STRING (*33*) and KEGG (*35*), but do not provide information about the nature of the relationships between genes, their protein products, or regulatory RNAs. **Supplementary Table 2C** lists the gene definitions for the members of the ketamine glutamate receptor sub-network. The Key for the symbols in this figure is shown in the caption of **Figure 3. Supplementary Table 2C** lists the gene definitions for the members of the ketamine glutamate receptor sub-network. The Key for the symbols is shown in the caption of **Figure 3**.

### GWAS disease risk SNPs and Hi-C loops discriminate 2 ketamine sub-networks

We hypothesized that psychiatric and social genotype-phenotype associations from GWAS may be useful in the functional characterization of the effects of ketamine in humans, in addition to published GWAS SNPs associated with dissociation and antidepressant measures in response to ketamine (*15, 37*). Because the majority of GWAS SNPs are located within intergenic and intragenic enhancers (*28, 30, 38*) we tested whether GWAS disease risk located within the ketamine glutamate receptor and ketamine neuroplasticity sub-networks were likely to be located within chromatin loops in neural cells (SK-N-SH, H1, A735 cell lines and postmortem brain). If the GWAS SNPs contained in the sub-networks were predicted to be causal in the appropriate human surrogate tissue types, either neural (SK-N-SH, H1) or astrocyte (A735) cells but not in a liver cell line (HepG2) or in a white blood cell using multiple machine learning algorithms (*39-45*), they were used to probe public Hi-C datasets using a bioanalytic method developed in our laboratory (*46*) **(Materials and Methods)**.

We found a total of 186 disease and pharmacogenomic association signals for SNPs predicted to be causal within the ketamine glutamate receptor sub-network and within the ketamine neuroplasticity sub-network. These 186 signals correspond to 108 non-overlapping SNPs and 78 SNPs were significantly associated with at least two traits. The 78 multi-trait SNPs were located either in the ketamine glutamate receptor sub-network or the ketamine neuroplasticity sub-network, but not in both sub-networks. Machine learning predicted that all of the sub-network SNPs were causal. The threshold for predicted causality required consensus of the numerical scores output from five different machine learning algorithms (*39-44*) and a deep learning software application (*45*) **(Materials and Methods)**. When compared to the scores generated from a random selection of ten GWAS SNPs for all human traits with *p*-values at 5E-08 or lower, every score for each of the GWAS SNPs contained within these two sub-networks was designated as casual **(Supplementary Table 4, Supplementary Table 5)**. Phenotype associations were carefully evaluated from source publications to eliminate those that were entirely self-reported or were ambiguous in terms of assignment to a coded psychiatric disorder (*37, 47*). The 186 GWAS signals associated with genes and regulatory RNAs that were members of the ketamine pharmacogenomic sub-networks identified in this study were annotated by disease risk, psychiatric subtype, enhancer, eQTL, Hi-C score (from both public data and our own adjustable bin mapping method) and co-localization with enhancer RNA. These 186 signals comprised 108 unique GWAS disease risk and pharmacogenomic SNPs predicted to be casual, including 78 SNPs with signals for multiple traits, exhibited clear genotype-phenotype association differences that discriminated the ketamine glutamate receptor sub-network from the ketamine neuroplasticity sub-network **(Table 1)**, although there was overlap for certain genetic associations including schizophrenia and smoking and recurrent depression **(Supplementary Table 4, Supplementary 5)**.

**Table 1.** Selected GWAS SNPs determined to be causal and annotated as enhancers located within (A) the ketamine glutamate receptor sub-network, and (B) the ketamine neuroplasticity sub-network. Data sources used for GWAS and SNP annotation: columns 1-5, the NHGRI-EBI GWAS catalogue (*37*), column 6, enhancer status determined using annotation from (*38, 48-60*), and column 7 using Hi-C data from (*46, 61, 62*). eQTL data support the results shown in column 7 *(63, 64)*. PMID in column 8 indicates PubMed identification number (*65*). All the GWAS SNPs analyzed can be found in **Supplementary Table 4** and **Supplementary Table 5**. *Significance increased by network boosting *(66)*.

| **A. The ketamine glutamate receptor sub-network** |  |  |  |  |  |  |  |
| --- | --- | --- | --- | --- | --- | --- | --- |
| **GWAS SNP** | **Populations** | **Gene(s)** | **Reported trait** | **Reported P-value** | **Enhancer, cell type** | **Hi-C score: Human brain** | **PMID** |
| rs1051730 | ASN, EUR | *CHRNA5, CHRNA3* | Smoking status | 6E-121 | Neuron | 1E-27: Nucleus accumbens | 30617275 |
| rs28681284 | ASN | *CHRNA5, CHRNA3* | Schizophrenia, chronic (F20) | 6E-13 | Neuron | 1E-27: Nucleus accumbens | 30285260 |
| rs2007044 | ASN | *CACNA1C* | Schizophrenia, chronic (F20) | 6E-20 | Bipolar neuron | 1E-10: Cingulate cortex | 29483656 |
| rs7192140 | EUR | *GRIN2A* | Smoking status | 2E-15 | Bipolar neuron | 1E-19: Frontal cortex | 30643251 |
| rs11647445 | EUR | *GRIN2A* | Bipolar I disorder (F31.0 - F31.64) | 1E-10 | Bipolar neuron | 1E-19: Frontal cortex | 31043756 |
| rs7893279 | ASN, EUR | *CACNB2* | Schizophrenia, chronic (F20) | 9E-14 | Neuron | 1E-10: Frontal cortex | 30285260 |
| rs111294930 | EUR | *LINC01470, GRIA1* | Schizophrenia, chronic (F20) | 9E-12 | Bipolar neuron | 1E-12: Cingulate cortex | 29483656 |
| rs9292918 | ASN | *HCN1* | Schizophrenia, chronic (F20) | 4E-11 | Bipolar neuron | 1E-10: Frontal cortex | 28991256 |
| rs62367520 | EUR | *HCN1* | Smoking status | 1E-10 | Bipolar neuron | 1E-10: Frontal cortex | 29283656 |
| **B. The ketamine neuroplasticity sub-network** |  |  |  |  |  |  |  |
| **GWAS SNP** | **Populations** | **Gene(s)** | **Reported trait** | **Reported P-value** | **Enhancer, cell type** | **Hi-C score: Human brain** | **PMID** |
| rs61902811 | EUR | *TMPRSS5, DRD2* | Recurrent depression (F33) | 4E-39 | Neuron | 1E-04: Hippocampus | 30718901 |
| rs4936277 | EUR | *TMPRSS5, DRD2* | Alcohol use disorder, alcohol dependence | 1E-13 | Bipolar neuron | 1.00E-06: Nucleus Accumbens | 29942085 |
| rs7227069 | EUR | *DCC* | Recurrent depression (F33) | 2E-28 | Neuron | 1E-10: Cingulate cortex | 30718901 |
| rs12967143 | EUR | *TCF4* | Recurrent depression (F33) | 2E-27 | Neuron | 1E-10: Frontal cortex | 30718901 |
| rs7932640 | EUR | *GRM5* | Recurrent depression (F33) | 3E-25 | Neuron | 1E-04: Cingulate cortex | 30718901 |
| rs139438618 | AFR | *SEMA3A* | Major depression and alcoholism | 2E-11 | Bipolar neuron | 1E-10: Nucleus Accumbens | 29071344 |
| rs775766 | EUR | *ROBO2* | Recurrent depression (F33) | 2E-08 | Neuron | 1E-14: Cingulate cortex | 29942085 |
| rs1400237 | EUR | *ROBO2* | Response to ketamine in depression | 8E-06* | Neuron | 1E-14: Cingulate cortex | 30552316 |

**Table 1A** shows that multiple GWAS disease risk SNPs located within the glutamate receptor sub-network can be annotated as enhancers associated with tobacco smoking status, chronic schizophrenia (ICD diagnostic code F20), and bipolar 1 disorder (ICD diagnostic codes F31.0 – F31.64). **Table 1B** shows that the ketamine neuroplasticity sub-network contains multiple GWAS disease risk SNPs that can be annotated as enhancers associated with recurrent depression (ICD code F33), alcoholism and response to ketamine. The results shown in **Table 1, Supplementary Table 4**, and **Supplementary Table 5** demonstrate that enhancer SNPs co-localize with a limited number of genes within each sub-network.

The ketamine glutamate receptor sub-network contains SNPs that occur with greatest frequency in association with *CHRNA3, CHRNA5, NCAM1, NOS3, CACN1AC, CACNB2, HCN1*, an intergenic region between *GRIA1 and LINC01470* and *GRIN2A*. The ketamine neuroplasticity sub-network contains SNPs that occur with greatest frequency in association with *TCF4, DCC*, an intergenic region between *DRD2* and *TRMPSS5*, and *GRM5*. These results demonstrate that mutational alteration of enhancer interactomes discriminate between the ketamine glutamate receptor sub-network (chronic schizophrenia, smoking status) and the ketamine neuroplasticity sub-network (recurrent unipolar depression, alcohol consumption).

### Ketamine enhancers are preferentially located in the cingulate cortex and the frontal cortex

The distribution of GWAS SNPs, enhancer, super-enhancer, and chromatin spatial contacts are concentrated within the cingulate cortex and the frontal cortex for both the glutamate receptor sub-network and the neuroplasticity sub-network. **Figure 6** shows the neuroanatomical distribution of the 108 GWAS SNPs located within Hi-C loops in either the ketamine glutamate receptor or ketamine neuroplasticity sub-networks based on the published literature (*46, 61, 62)*. These results, combined with correlative mapping of gene expression within these sub-networks based on data from the human brain atlas of the Allen Brain Science Institute (*67*) and functional neuroimaging results from 24 studies of ketamine’s rapid antidepressant response **(Supplementary Table 1)**, support research demonstrating that cingulate cortex and frontal cortex are primary sites where the drug exerts its pharmacodynamic effects (*68, 69*).

**Fig. 6.**
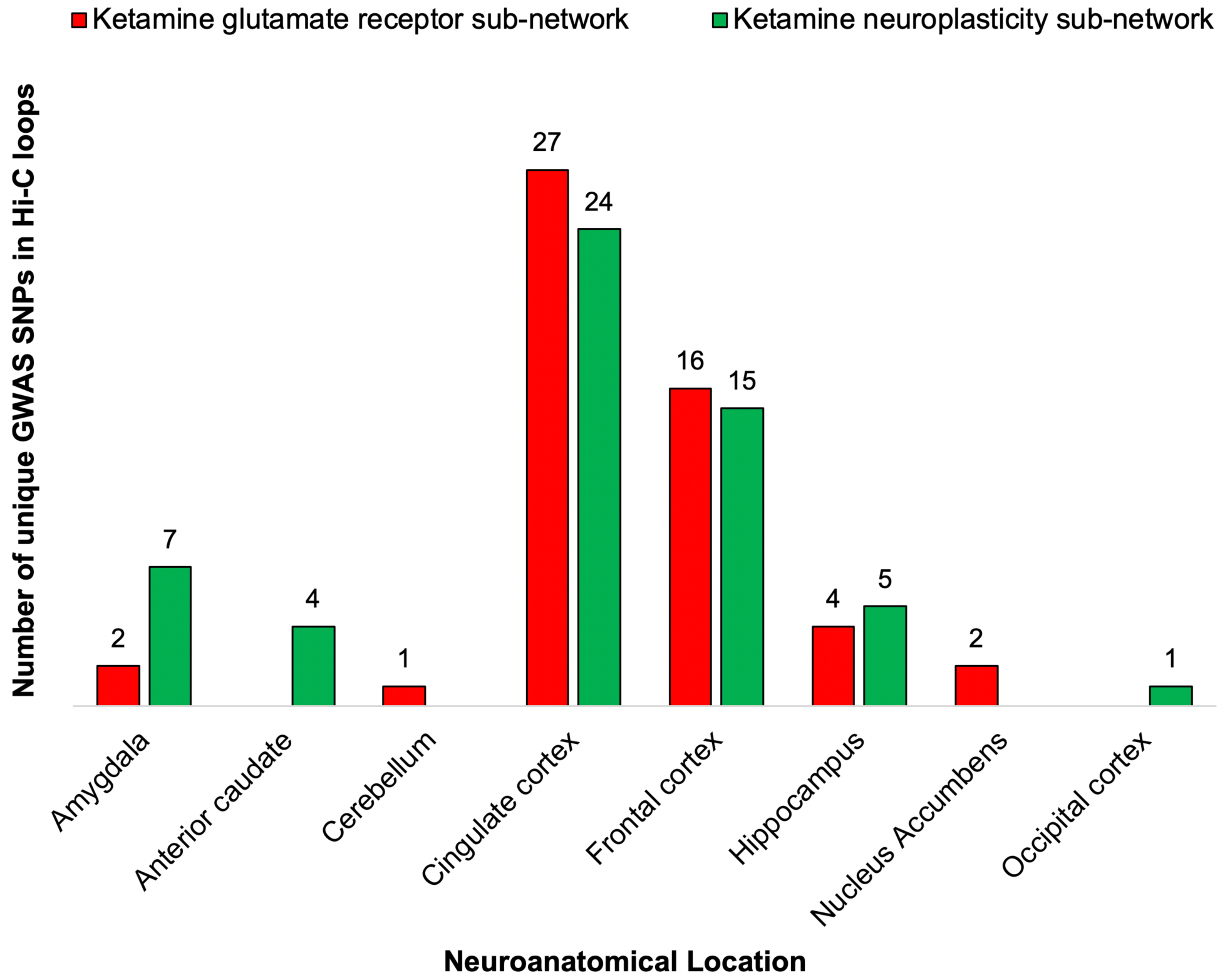
The neuroanatomical distribution of 108 non-overlapping GWAS disease risk and pharmacogenomic response SNPs located within Hi-C chromatin loops in the ketamine glutamate receptor sub-network (red) and the ketamine neuroplasticity sub-network (green). Details provided in **Supplementary Tables 4 and 5**.

Evidence in support of the ketamine pharmacogenomic network identified here is demonstrable by comparing the neuroanatomical distribution of gene expression data within the ketamine sub-networks with the localization results from a consensus brain-map showing which brain regions are first impacted by ketamine obtained from 24 neuroimaging studies **(Supplementary Table 1)**. The consensus map emphasizes the anterior cingulate cortex (ACC), dorsolateral and dorsomedial prefrontal cortex (PFC), and the supplementary motor area (SMA) as consistently the first human brain regions to be activated by the drug, indicated by the dark red spheres in **Supplementary Figure 1A**. However, other CNS regions have been reported in neuroimaging studies to be rapidly impacted by ketamine in humans following administration of the drug. These are shown in black in **Supplementary Figure 1A** but did not comprise the clear majority of brain regions reported to be first impacted by ketamine in the neuroimaging studies that we examined during our research. To serve as controls, we chose adjacent human brain regions not impacted by ketamine in the neuroimaging studies, including the corpus callosum (CC), occipital cortex (OC) and somatosensory cortex (SS). We were limited by available sources of gene expression data from postmortem human brain but did obtain reliable and non-conflicting data on 100 of the 107 genes in the ketamine network (*64, 67*). Also, the RNA sequencing (RNA-seq) results were not as well localized as the neuroimaging data, so this analysis was limited to obtaining RNA-seq data labeled as originating from anterior cingulate cortex (ACC), prefrontal cortex (PFC), corpus callosum (CC), somatosensory cortex (SS), and occipital cortex (OC). As shown in **Supplementary Figure 1B**, genes in the ketamine network are expressed at significantly higher levels in the ACC and PFC than in the neighboring CC, SS and OC, where there is no evidence that ketamine exerts rapid antidepressant effects. The ACC is part of the cingulate cortex, and the PFC is part of the frontal cortex, regions that are shown in **Table 1, Supplementary Figure 2, Supplementary Table 4** and **Supplementary Table 5**.

Publicly available *in situ* hybridization data of *GRIN2B, GRIN1* and *GLRB* mRNA in postmortem human brain **(Supplementary Figure 2)** (*67*) shows that expression of these three key genes located in the ketamine glutamate receptor network is widespread in the telencephalon, including amygdala, anterior caudate, anterior cingulate cortex, frontal cortex, globus pallidus, hippocampus, prefrontal cortex, nucleus accumbens and putamen. However, there are differences in their neuroanatomical distribution. For example, *GLRB* shows circumscribed expression in layer VI of cortex, whereas *GRIN2B* and *GRIN1* are expressed in the dentate gyrus of the hippocampal formation.

Whole genome, Hi-C data mapping performed using SNPs as data probes, the probes including SNPs contained within the ketamine sub-networks and obtained from the GWAS catalog *(37)*, including those which has been significantly associated with disease risk and ketamine antidepressant response variation and dissociation (*15*). These results validated pathway analysis and demonstrated both *cis*- and *trans*-interactions with other members of the ketamine pharmacogenomic pathway within human neurons **(Figure 7, Figure 8) (Materials and Methods)**. These spatial contacts are significantly enriched for association with specific superenhancers from cingulate cortex and frontal cortex (*data not shown*). **Figure 7A** shows a whole genome plot that is the key for understanding the gene-gene interactions shown in **Figure 7B-7G**, with each individual *trans*-interaction labeled using a different color (see Key). **Figure 7B** shows Hi-C contacts between *RASGRF2*, a gene associated with synaptic plasticity and alcoholism *(70)*, with the co-localized nicotinic receptor genes *CHRNA3* and *CHRNA5* that contain SNPs significantly associated with smoking status in GWAS *(37)*. **Figure 7B** shows *trans*-interactions between the *ROBO2* gene, which contains a number of SNPs associated with both unipolar depression and dissociative and antidepressant responses to ketamine in GWAS *(15,37)*, and both the *GRIN2B* gene and the *ATF7IP* gene. The *ATF7IP* gene encodes a chromatin remodeling protein responsible for HUSH-mediated heterochromatin formation and gene silencing as part of the stabilization of the SETDB1 complex, required for methylation of histone 3 lysine 9 (H3K9me3) during neuronal differentiation *(71)*. The *TCF4* gene, encodes a transcription factor that regulates neuronal differentiation *(15, 37)*, and harbors multiple GWAS SNPs significantly associated with recurrent depression **(Supplementary Table 4)**.

**Fig. 7.**
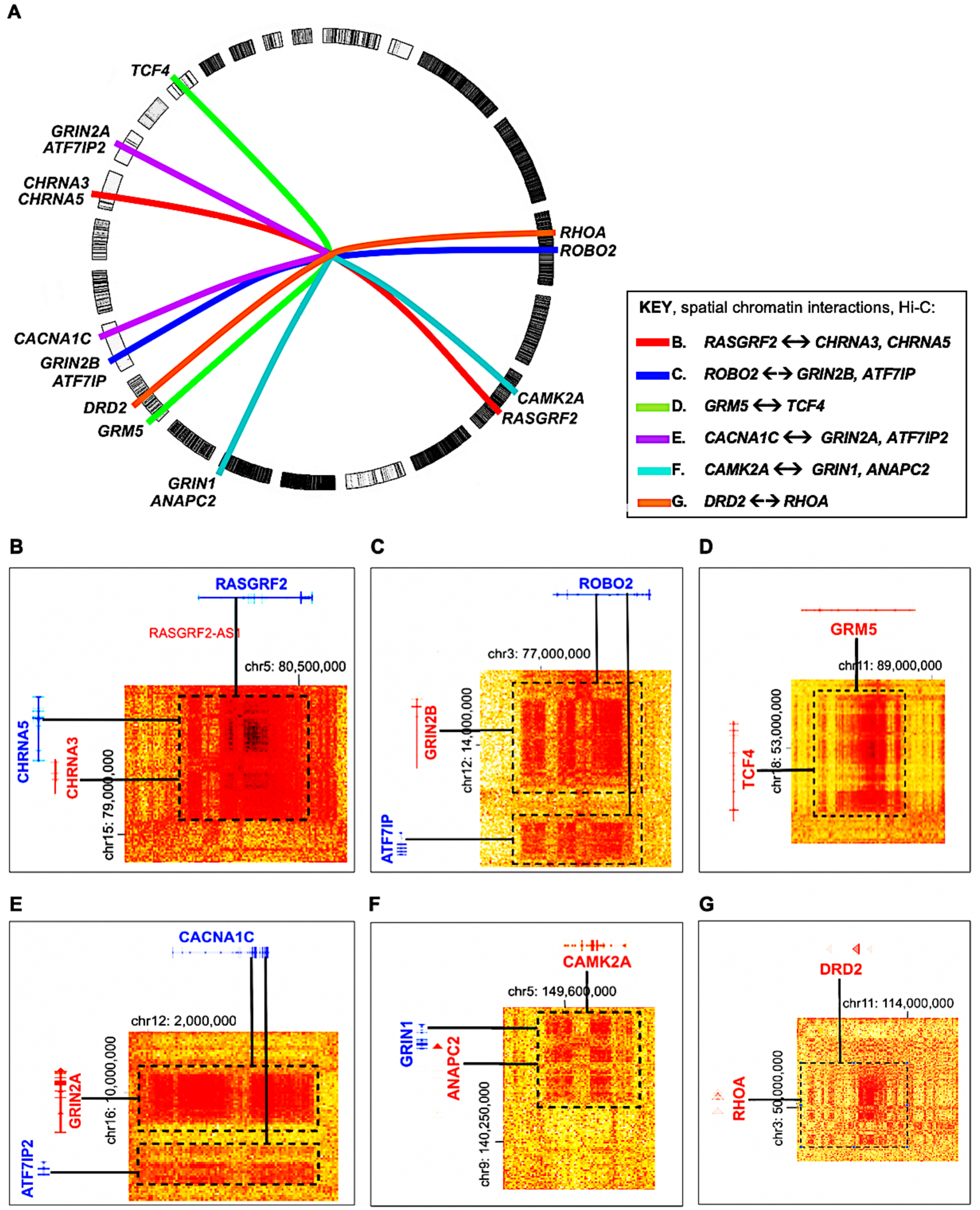
Example of the *trans*-interactions of ketamine pharmacogenomic network SNPs. The intensity heat maps demonstrate individual spatial Hi-C contacts between genes in human neurons, including **(A)** *CACNA1C* and *GRIN2A* (blue line); **(C)** *RASGRF2* and *CHRNA3 CHRNA5* (orange line); **(D)** *CAMK2A* and *GRIN1, ANPAC2* (red line); **(E)** *DRD2* and *RHOA* (light blue line); **(F)** *GRM5* and *TCF4* (purple line); (G) *ROBO2* and *GRIN2B, ATF7IP* (green line); **(B)** Whole genome plot serves as key showing individual gene-gene interactions by color with the corresponding colored line located next to letters in Figure 7A, C-G. Images from Hi-C data visualization using the HiGlass software *(74)*, and coordinates are based on human genome build hg19 of the UCSC browser (*75*).

**Fig. 8.**
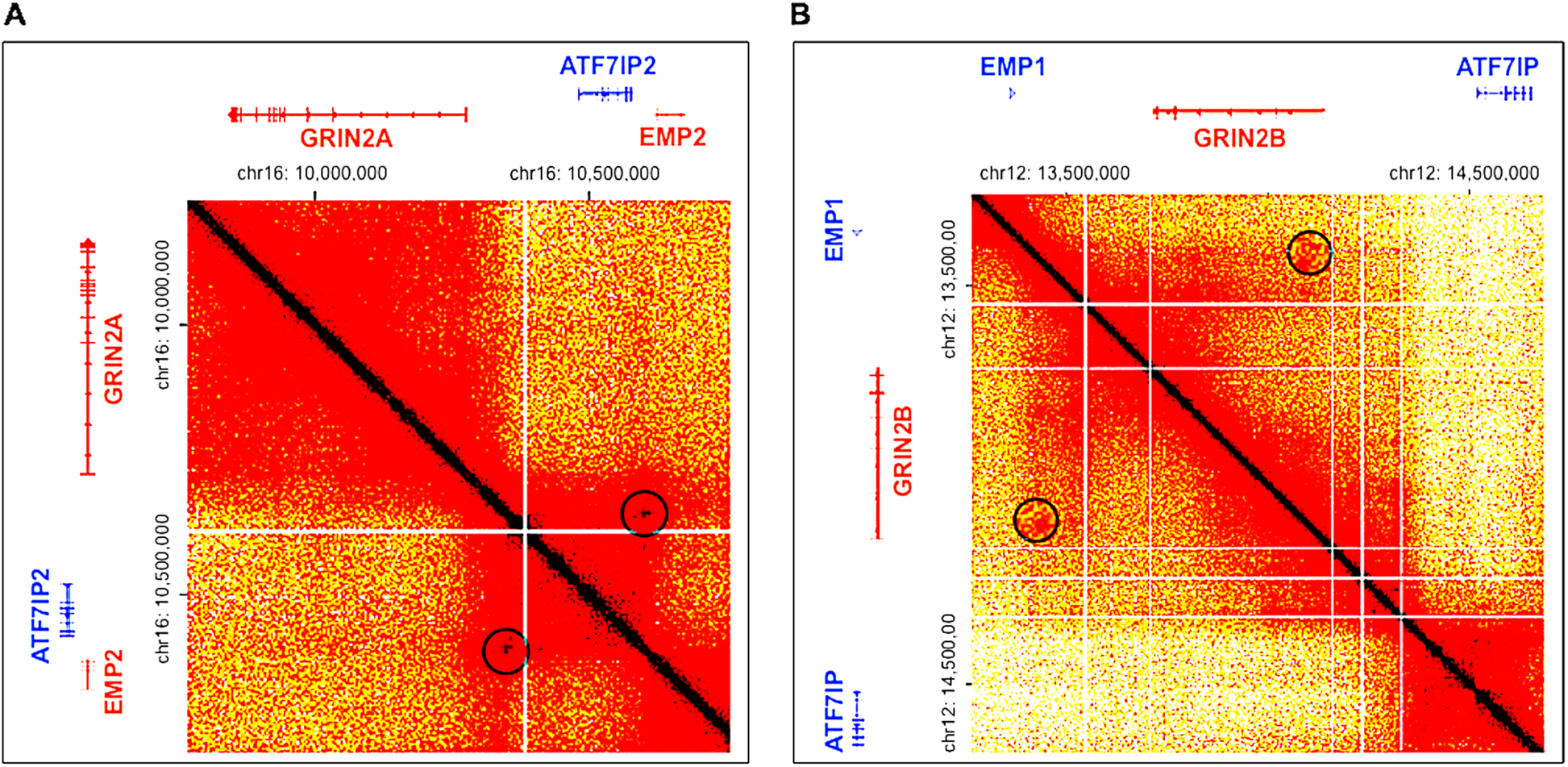
Organization of the Hi-C loci containing the glutamate receptor genes *GRIN2A* and *GRIN2B* in human glutamatergic neurons. **(A)** Hi-C matrix showing the co-localized genes *GRIN2A, ATF7IP2* and *EMP2* **(B)** Hi-C matrix showing the co-localized genes *GRIN2B, ATF7IP* and *EMP1*. Circles indicate possible loop domains magnified by 2X. Images from Hi-C data visualization using the HiGlass software *(74)*, and coordinates are based on human genome build hg19 of the UCSC browser (*75*).

**Figure 7D** demonstrates a spatial contact between *TCF4* and the *GRM5* gene, which encodes a member of the glutamate metabotropic receptor family and contains enhancers significantly associated with depression in GWAS **(Supplementary Table 4)** *(37)*. **Figure 7E** shows a Hi-C map of interactions between *CACNA1C* and the *GRIN2A* and the ATF7IP2 genes. In **Figure 7C**, Hi-C spatial contacts obtained from human glutamatergic neurons shows *trans*-interactions between the *CAMK2A* gene located on chromosome 5 with the genes *GRIN1* and *ANAPC2* located on chromosome 9. The ANAPC2 protein is part of a complex that controls the formation of synaptic vesicle clustering at the active zone to the presynaptic membrane in postmitotic neurons, and this complex also degrades NEUROD2 as a primary component of pre-synaptic differentiation during neuronal differentiation *(72)*. **Figure 7G** shows spatial contacts in neurons between the *DRD2* gene and the *RHOA* gene, which encodes a signaling protein that regulates the cytoskeleton during synaptic transmission in neurons *(73)*.

**Figure 8** compares the local *cis*-interactions and organization of the loci containing the *GRIN2A* and *GRIN2B* genes in human glutamatergic neurons. In both cases, related genes are organized as triplets on different chromosomes, with the *GRIN2A, ATF7IP2* and *EMP2* genes on chromosome 16, and the *GRIN2B, ATFIP* and *EMP1* genes on chromosome 12. It is not known whether the co-localization of these 3 related genes is a coincidental, possibly a consequence of a conserved genomic duplication occurring during evolution and/or functional relatedness. **Figure 8** shows putative chromatin loops (magnified circles).

## Discussion

### The ketamine pharmacogenomic sub-networks

We have identified a ketamine pharmacogenomic network in the human brain using multi-scale biological data analysis. Our approach used multiple, reinforcing data sources and algorithms, as well as results spanning different levels of resolution, ranging from disease risk genotype-phenotype associations obtained from GWAS to functional neuroimaging of rapid antidepressant response in the human brain following ketamine administration. Special emphasis has been placed on discretization of the enhancer interactome and chromatin interactions in the context of pharmacogenomic efficacy and adverse events. Results from all of the methods applied at different scales of resolution produced consistent findings, demonstrating that this novel strategy provides a useful application for data-driven bioinformatics discovery of psychotropic drug mechanisms for future applications **(Figure 1)**. The research reported here also provides an *in-silico* design framework for subsequent study of pharmacogenomic networks in other therapeutic categories.

This research emphasizes the role of the regulatory genome, including enhancer and superenhancer-based interactomes, as an approach that provides insight into pharmacogenomic network mechanisms (*22-25*). It differs from pathway modeling methods in which altered protein folding based on missense codon variants and fixed signaling pathways serve as the foundation for the interpretation of the molecular substrate of ketamine response in humans. For example, we found no evidence for the inclusion of SNPs within proposed key signaling pathways in the ketamine network such as *AKT1, MTOR*, or *ERK1*, or participation of the inflammasome in this network (NLRP3 pathway) (*10, 17, 76, 77*). Although we do not find inflammatory genes in this ketamine network, the drug has shown promise as an investigational therapy for some inflammatory disorders *(8)*. We did find evidence supporting the inclusion of the pharmacokinetic genes *CYP2B6* and *CYP3A4*, in addition to *ESR1* and *GRIA1, GRIA4* within the ketamine network in human brain consistent with recent studies (*12, 78*), although more study is needed to show how proteins encoded by these genes in the brain contribute to ketamine response in humans.

One possible limitation of this research is that it relies, in part, on public and commercial databases that may contain erroneous information including uncorrected confirmation bias in published literature, deprecated primary source material, and results obtained using flawed statistical methods and/or methods based on spurious assumptions. To mitigate error associated with reliance on a single data type, the approach in this study used multiple, related data sources and algorithms, spanning different levels of resolution ranging from disease risk genotype-phenotype associations obtained from GWAS to functional neuroimaging of rapid antidepressant response in the human brain following ketamine administration. Similar methods have been used to combine results from GWAS and neuroimaging data *(79)*.

### The three-composite ketamine pharmacogenomic sub-networks

This research suggests that a different pharmacogenomic sub-network may mediate antidepressant effects and the dissociative and psychotomimetic effects of ketamine. Thus, the function of one sub-network is characterized by neurogenesis and neuroplasticity, as previously shown as the mechanism of action for other antidepressant medications (*9, 80, 81*). It is tempting to speculate that while ketamine may not directly cause neurogenesis in its role as an antidepressant in human brain, it may stimulate neurogenic transcription leading to renewal of intrinsic self-efficacy, a personality trait often lost in depression (*82*). In contrast, the sub-network containing genes that encode glutamate receptor proteins such as GRIN, GRIN3A and GRIN2B not only contains the largest number of genes, but SNPs within enhancers of this sub-network have been significantly associated with dissociative disorders such as schizophrenia and bipolar disorder, and postoperative delirium, vomiting, and nausea adverse events associated with ketamine administration. Finally, enhancer SNPs in the glutamate receptor sub-network versus the neuroplasticity sub-network stratify by ketamine efficacy with reduced dissociation versus increased dissociation in response to ketamine, respectively (*15*).

The ketamine pharmacogenomic network is preferentially localized in the anterior limb of the behavioral antidepressant connectivity network, including cingulate cortex and frontal cortex in the human brain. This connectivity network was confirmed using an array of different data types, including enhancer, superenhancer, eQTL and Hi-C scoring of chromatin loops for annotation of GWAS SNPs, expression mapping, functional neuroimaging studies, and examination of the *cis*- and *trans*-interactions of ketamine pharmacogenomic SNPs.

There is a need to better understand mechanisms through which ketamine and its analogues act in the human central nervous system and to uncouple the psychotomimetic adverse effects of this emerging class of promising therapeutics from mechanisms that mediate antidepressant action. Ketamine’s action as an anesthetic, analgesic and sedative may be a consequence of the partial antagonism of NMDAR-mediated pain transmission, blocking central sensitization of ascending pain afferents from the dorsal horn. Thus, through dissociation, the patient no longer pays attention to ascending pain stimuli, detached from the conscious mind, in a dreamlike state often accompanied by memory impairment (*83, 84*).

### Genotype-phenotype annotation of different ketamine sub-networks

One challenge is to understand why certain phenotypes cluster within a specific ketamine pharmacogenomic sub-network. In the glutamate receptor sub-network of the larger ketamine pharmacogenomic network, significant genotype-phenotype associations that include smoking impacting the same genes as schizophrenia **(Table 1A and Supplementary Table 4)**. In the ketamine neuroplasticity sub-network, significant enhancer annotations included unipolar depression, depressive symptoms and alcoholism. This lends credence to our hypothesis that mutational disruption of the enhancer interactome within a drug network may provide insight into pharmacogenomic response stratification.

There is substantial evidence that patients diagnosed with schizophrenia are more likely to be heavy smokers than are healthy controls (*85-87*) but cause and effect are not understood. Emerging research demonstrates that nicotine induces *CYP2B6* gene expression by two to three orders of magnitude in the human brain but not in the human liver (*14*). Because the cytochrome P450 family 2 sub-member 6 enzyme encoded by the *CYP2B6* gene is rate-limiting in the metabolism of ketamine and its enantiomers (*88*), patients suffering from one of the spectrum of dissociative disorders related to schizophrenia may use nicotine for self-medication (*85*); however this hypothesis remains speculative. While exon variants in the *CYP2B6* gene may differentially alter splicing but not protein structure or function (*89*), mutations that alter *CYP2B6* mRNA splicing in the human brain contribute significantly to ketamine response variation in humans (*90, 91*). Previous research and results from this study show that estrogen regulation of *CYP2B6* gene expression can modulate differential response to ketamine in females (biological sex) **(Figure 5)** (*92*). These results indicate that dose optimization is critically dependent on a patient’s endogenous drug profile, biological sex, and age *(12, 91, 92)*.

### Translation to clinical practice

These results support the conclusion that variation in genomic regulatory networks can be translated into actionable drug selection and dose optimization in patients. This study demonstrates the potential of multi-scale biological data analysis for the *in-silico* prediction of drug response and serious adverse event profiles in patients based on normal variation in the non-coding enhancer interactome and the discretization of drug-disease networks based on common genotype-phenotype associations. Thus, the antidepressant and dissociative effects of ketamine and related glutamatergic modulators, including dextromethorphan, D-cycloserine, raspastinel and sarcosine, may be uncoupled based on the patient’s genetic risk profile. In this manner, knowledge of the subordinate mechanistic sub-networks of ketamine and its enantiomers explain previously unrecognized mechanisms of action of this drug class, identifying additional targets for drug repurposing. This strategy also suggests possible clinically relevant indications for ketamine that extend beyond depression, surgical anesthesia, sedation, and acute analgesia. These include psychiatric and also stress-related disorders defined by similar mechanisms of neural injury and repair, including posttraumatic stress disorder, bipolar 1 disorder, fibromyalgia, peripheral neuropathy, inflammatory bowel disease, and global chronic pain. The clinical utility of this approach requires confirmation in preclinical and clinical studies.

## Materials and Methods

### Selection of ketamine-response genes

The ketamine response workflow is based on the pharmacoepigenomics informatics pipeline (PIP) *(22-25)*. Input genes to the data analysis pipeline first included those genes that encode proteins and constituents of macromolecular protein complexes obtained from past studies of the binding affinity of (*R, S*)-ketamine, *R*-ketamine and *S*-ketamine performed in microsomal and tissue preparations in rodents and humans. Genes were selected based on whether their protein products, either alone or as part of a larger complex, bound ketamine enantiomers at high affinity ranging from 0.1 – 100 μM. To identify proteins and protein complexes, we used a search string from January 1975-October 2019 that included a Boolean search string with the words, “ketamine’”, “*(RS)-*ketamine”, “*(R)*-ketamine”, “*(S)*-ketamine”, “binding affinity”, “K_i_” and “IC_50”,_ “brain”, and “microsomes” – this search string produced 18 publications **(Supplementary Table 1)**. This gene set included *CACNA1C, CHRM2, CHRNA3, CHRNA5, CHRNA7, CNR1, DRD2, ESR1, GLRB, GRIA1, GRIA4, GRIN1, GRIN2A, GRIN2B, GRIN2C, GRIN2D, GRIN3A, GRIN3B, GRM5, HCN1, HTR1A, HTR1B, HTR2A, HTR3A, OPRK1, OPRM1, SIGMAR1, SLC6A2 and SLC6A3* (see also **Supplementary Table 3**.)

Further inputs identified candidate pharmacodynamic and pharmacokinetic genes obtained from published literature and drug databases, including DrugBank *(93)*, Ingenuity Pathway Analysis™ (IPA™; Qiagen GmBH) *(32)*, the Kyoto Encyclopedia of Genes and Genomes (KEGG) *(35)*, and DrugCentral *(94)*. This subset included genes not listed in the above set of candidate genes, but excluded pharmacodynamic targets that had not been shown to bind ketamine and its enantiomers at high affinity. This subset included genes and regulatory RNAs: *ACHE, AKT1, ANAPC2, AR, ARC, ARNT, ASCL1, ATF7IP, ATF7IP2, ATP1A1, BDNF, BDNF-AS, BORCS7, CACNA1C, CACNB1, CACNB2, CACNG2, CAMK2A, CDKN1A, CREM, CUX2, CYP2A6, CYP2B6, CYP2C9, CYPC19, CYP3A4, DCC, DLG3, DLG4, DNMT1, DOCK10, EDN2, EEF2, EEF2K, EHTM1, ENSG00000225960, ENSG00000251574, ERK1, ERK2, ESR1, FMR1, GABBR1, GABRA2, GABRA5, GAD1, GDAP1L1, GFAP, GLRA1, GLRA2, GRIP1, HDAC5, HK1, HMOX1, HOMER1, KLF6, LAMTOR1, LEP, LIN7C, MAPK1, MAPK8, MBD1, MEF2D, MPHOSPH8, MTOR, MTRNR2L2, MYO6, MYT1L, NMB, NCAM1, NEUROD1, NEUROD2, NLRP3, NMB, NOS1, NOS2, NOS3, NQO1, NR4A1, NSMF, NTRK2, PGBD1, PTGS2, PVALB, RAC1, RASGRF2, RHOA, ROBO2, RPTOR, SEC11A, SEMA3A, SETDB1, SHANK1, SHANK2, SHANK3, SLC6A9, SLC22A15, SLIT1, SLIT2, SNAP25, SYN1, SYN2, SYT3, TASOR, TBR1, TBX21, TCERG1, TCF4, TMPRSS5, TMPRRS6, TOGARAM2, TRIM26, TRIM28, ZNF274, ZNF592* and *ZNF717*.

Criteria for further selection of candidate genes was 2-fold. First, we determined whether the gene or regulatory RNA was expressed in human brain regions rapidly activated by ketamine enantiomers as determined by functional neuroimaging. The neuroanatomical distribution of the expression of genes in the ketamine network and its sub-networks were matched to a consensus map of brain regions rapidly activated by ketamine, based on the results of 24 neuroimaging studies in healthy controls and patients diagnosed with unipolar and bipolar depression **(Supplementary Table 1)**. Second, we assessed whether this gene set formed a significantly interconnected pathway in human brain using neural cell- and tissue-specific filtering. Results were generated using IPA™ (Qiagen GmBH) and verified using the STRING protein-protein interaction network database *(33)*. Independent software applications for the *post-hoc* bioinformatics analysis included enrichment analysis using Gene Ontology’s Panther web-based portal *(34)* and the gene expression module of IPA™ to determine disease risk based on published literature *(32)*.

### Gene set optimization, pathway analysis and significant upstream molecular regulators

Following reconstruction of the larger ketamine spatial network, it was deconstructed into its associative mechanistic sub-networks using an executable gene set optimization engine that iteratively compares gene function with efficacy, adverse events and other known features associated with ketamine response **(Figure 9)**. The iterative mechanistic sub-network profiler comprises executable code that iteratively re-organizes interconnected gene sets within the drug spatial network until the most significant associations with the various mechanisms of action of the psychotropic drug are reached, based on multiple databases including Gene Ontology *(34)*.

**Figure 9.**
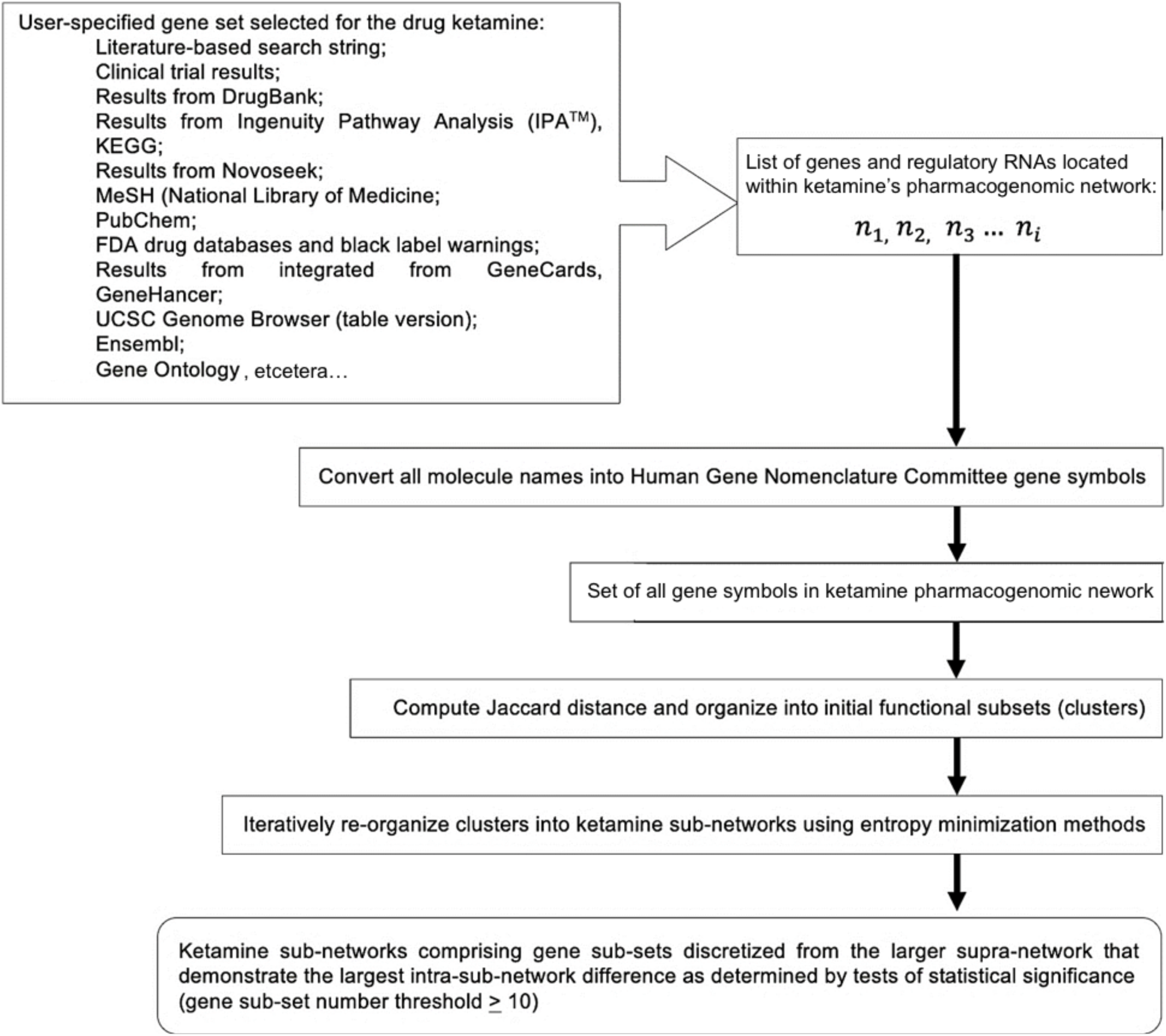
Iterative gene set optimization workflow. This iterative gene set optimization executable differs from gene set enrichment methods, by not only combining different mathematical methods, but also not acting in a hierarchal manner ranking genes as in threshold-dependent methods until the output, and this iterative gene set optimization does not rely on comparisons of experimental results, such as in whole-distribution tests. Instead, the iterative gene set optimization groups genes or long noncoding RNAs using the Jaccard distance to first measure the similarity between two genes or long noncoding RNAs based on the dissimilarity of user-selected terms, where the Jaccard distance is as the ratio of the size of the symmetric difference *Gene A* Δ*Gene B* = *A*⋂*B* − *A*⋃*B* to the union. This is extensible into clusters of related dissimilar gene names. A drug network server then automatically sorts these sets, or using user-defined numbers of clusters, into subsets of clustered subsets of functionally related genes using a minimal entropy sorting algorithm, such as the COOLCAT algorithm *(94)*. Following gene subset optimization using entropy minimization, the pharmacogenomic spatial network identification system may employ manual curation to assign efficacy, adverse event or functional mechanistic sub-networks based on known attributes of the drug’s mechanism of actions under consideration.

Independent pathway analysis software applications were used for the *post-hoc* bioinformatics analysis including the gene expression module of IPA™ *(32)* and KEGG *(35)* to determine disease risk, based on results extracted from the published literature. In addition, ketamine sub-network genes were annotated using data from the EBI-NHGRI GWAS catalog *(37)* to assess the potential of sub-network function through the identification of significant SNP-trait associations for every gene in each sub-network. We hypothesized that mutations within gene subsets that comprise each sub-network, as exemplified by SNP-trait associations from GWAS, were determined to be relevant to ketamine efficacy and adverse events and, as such, might provide further insight into the normal, unimpaired function of the various sub-networks.

### Selection of candidate ketamine response SNPs

Our pharmaco-informatics platform processes data with minimal human intervention to avoid in-house bias. Human SNPs were selected and processed using a revised version of the bioinformatics pipeline that has been described in detail in previous publications *(22-25)*. For the workflow used in this study, GWAS SNPs were used as reported. These SNPs included all variants from GWAS related to genes within the larger ketamine pharmacogenomic network including genes located in the composite sub-networks. These included both SNP signals significantly associated with disease risk and those associated with ketamine antidepressant response efficacy and dissociation, as well as other ketamine network and sub-network-associated SNPs from the published literature and clinical trials *(8, 37)*. The SNP filter judged candidate ketamine network SNPs for putative causality using multiple, redundant machine learning algorithms *(39-45)*. The 65,535 SNPs contained in the EBI-NHGRI GWAS catalog *(37)* were evaluated for the 110-putative ketamine-response genes and regulatory RNAs contained in our sub-networks. Phenotypes that were defined by self-reporting, such as educational attainment, mathematical ability and intelligence, were not included. Phenotypes, including insomnia and neuroticism, were also not included as there are various interpretations of these psychiatric surrogate associations. Since a plethora of GWAS SNPs associated with schizophrenia and depression were found to be primary determinants of differences between the ketamine glutamate receptor sub-network and the ketamine neuroplasticity sub-network, we undertook to provide greater clarification of these broad-spectrum phenotypes into subtypes. Phenotypes including smoking status, alcohol and substance abuse, nausea, vomiting and migraine were included based on their relevance to the psychological features of patients or the mechanisms of action of ketamine and its enantiomers. The genotype-phenotype association annotations were examined for determination of differences between the ketamine glutamate receptor sub-network and the ketamine neuroplasticity sub-network.

Numerical scores from each algorithm are generated for each GWAS SNP and only in cases where each output scored the SNP as predicted to be causal in SK-N-SH cells and H1 cells, but not in HepG2 cells and PBMCs, were the SNPs retained for further analysis. These numerical scores for every SNP are shown in **Supplementary Table 4** and **Supplementary Table 5**. The score of every predicted causal SNP was independently tested to determine if it differed significantly from the scores generated using 10 randomly selected GWAS SNPs for all human traits at *p*≤5E-08 listed in the EBI-NHGRI GWAS catalogue *(34)* using ANOVA. Only when the SNP met this criterion of significance, was it selected for further analysis. An example for the SNP rs12967143-G, an intragenic enhancer within the *TCF4* gene, is shown in **Figure 10**.

**Figure 10.**
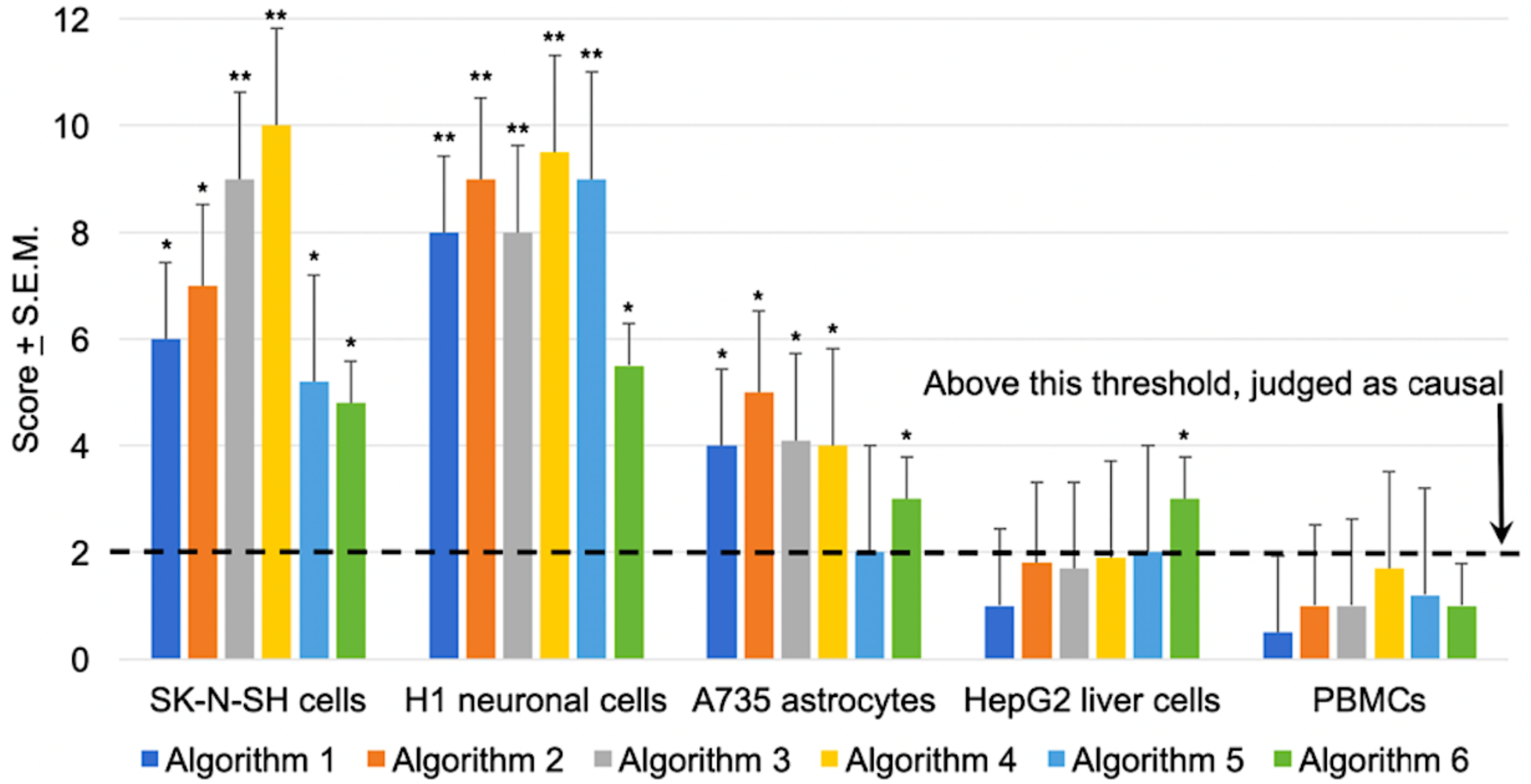
Results of significance testing of the SNP rs12967143-G, an intragenic enhancer located in the *TCF4* gene, versus other GWAS SNPs as described using the numerical output from the machine learning algorithms used in the analysis. \**p*≤0.05; ** *p*≤0.01.

### Annotation of enhancer-promoter pairs and chromatin contacts in neural cell lines and brain

GWAS SNPs within the 2 different ketamine sub-networks were annotated by disease risk, psychiatric subtype, enhancer, promoter, superenhancer and enhancer RNA co-localization, enhancer promoter interactions determined by Hi-C chromatin interactions using public data sources. We used an advanced version of an adjustable bin mapping method (formerly called “Hi-C compiling with Genomic and Regulatory Elements and Empirical Normalization”(“H-GREEN”)) method in which candidate ketamine response SNPs judged as causal using machine learning were used to probe public sources of Hi-C data obtained from SK-N-SH, H1, and A735 cells. This approach was used to verify public data on the localization of enhancer-promoter pairs. The method employs non-fixed, adjustable binning analytic software, which provides much higher resolution for point-to-point mapping in squared human genome space than other methods. The adjustable bin mapping method maps ketamine SNPs to determine the distribution of TADs altered by various psychotropic drugs within the chromatin interactome and the *cis*- and *trans*-interactions of TADs containing pharmacogenomic SNPs for ketamine that were assessed in neuronal, astrocyte, hepatocyte cell lines and peripheral blood mononuclear cells.

To map genomic element contacts, this chromatin interaction method obtains a set of genomic loci and segments the set into bins of varying sizes. The method selects 2 sets of bins corresponding to the home TAD of the pharmacogenomic SNP and one of its contact TADs (e.g., a first set of bins corresponding to chromosome 1 and a second set of bins corresponding to chromosome 8) and places them in an *n* x *m* matrix (a squared genome area) to generate a set of bin pairs. Accordingly, the squared genome area may be of variable size and shape. In some run versions, both sets of bins are the same (e.g., each corresponding to chromosome 1). The chromatin interaction system identifies pairs of locations corresponding to paired end reads or other spatially interacting locations with bin pairs that contain them, i.e. wherein one of the bins contains one locus and the other bin contains the other locus, using a binary search tree. Then an interaction frequency is generated for each bin pair based on the genomic element contacts within the corresponding bin pair. The interaction frequencies are normalized per the density of pairwise contacts as a function of genomic distance within each bin pair. More specifically, the density of pairwise contacts as a function of genomic distance is generated using a normalized density function.

For a bin pair containing origin and target contact TADs, the density function is integrated over the squared genome area of the bin pair to determine an expected density for the bin pair. The expected density is then compared to the actual density for the bin pair (i.e., the number of pairwise contacts within the squared genome area of the bin pair) using the statistical test of Benjamini-Hochberg false discovery rate *(96)*. These generate a collection of enriched and depleted chromatin contacts in a manner adjusted for distance (and other features as appropriate), on a local and genomewide basis. The chromatin interaction system then provides indications of the bin pairs having, for example, enriched or depleted contacts for display on a user interface. By using variable bin sizes, the chromatin interaction system enables adjustable Hi-C bin mapping, regardless of bin length.

### Neuroanatomical localization of enhancer sub-networks

First, comparison was made between the expression patterns of genes within the ketamine sub-networks with a consensus heatmap of the drug’s rapid antidepressant response map derived from functional neuroimaging, as previously described **(Supplementary Figure 1)**. Second, we evaluated the 108 non-overlapping GWAS SNPs annotated for their putative regulatory function as enhancers, promoters, superenhancers and splice variants, and then used a variety of sources of Hi-C data for neuroanatomical assignment to human brain regions including the amygdala, anterior caudate, cerebellum, cingulate cortex, frontal cortex, hippocampus, nucleus accumbens and the occipital cortex from Hi-C data *(46, 61, 62, 74)*.

#### Whole genome, Hi-C mapping performed using disease risk and pharmacogenomic SNPs

In addition to the adjustable bin mapping method developed in our laboratory, we used a local instance of the HiGlass software for rapid identification of the *cis*- and *trans*-interactions of all of the 108 non-overlapping GWAS SNPs shown in **Supplementary Table 4** and **Supplementary Table 5** *(74)*. This client-server software converts single resolution 1D linear human genome into multi-resolution formats that can be interactively searched in Hi-C from neuronal cells and tissues, and other sources, for spatial chromatin contacts. This software was used to generate a set of spatial interactions for the ketamine glutamate receptor sub-network and the ketamine neuroplasticity sub-network. In excitatory human glutamate neurons, the following have both *cis*- and *trans*-interactions with all members of this gene set: *ATFIP2, CACNA1C, CACNG2, CHNRNA3, DRD2, GRIN1, GRIN2A, GRIN2B, NOS1 and SETDB1*, compared to other tissues or cell lines. In neuronal cell lines, the following genes had both *cis*- and *trans*-interactions with each other: *TCF4, DCC*, and *GRM5*, compared to other tissues or cell lines. Examples of gene pair *trans* interactions within and between the sub-networks is shown in **Figure 7**, and local *cis* interactions are shown for the *GRIN2A* and *GRIN2B* loci in **Figure 8**.

## SUPPLEMENT

**Supplementary Table 1:**
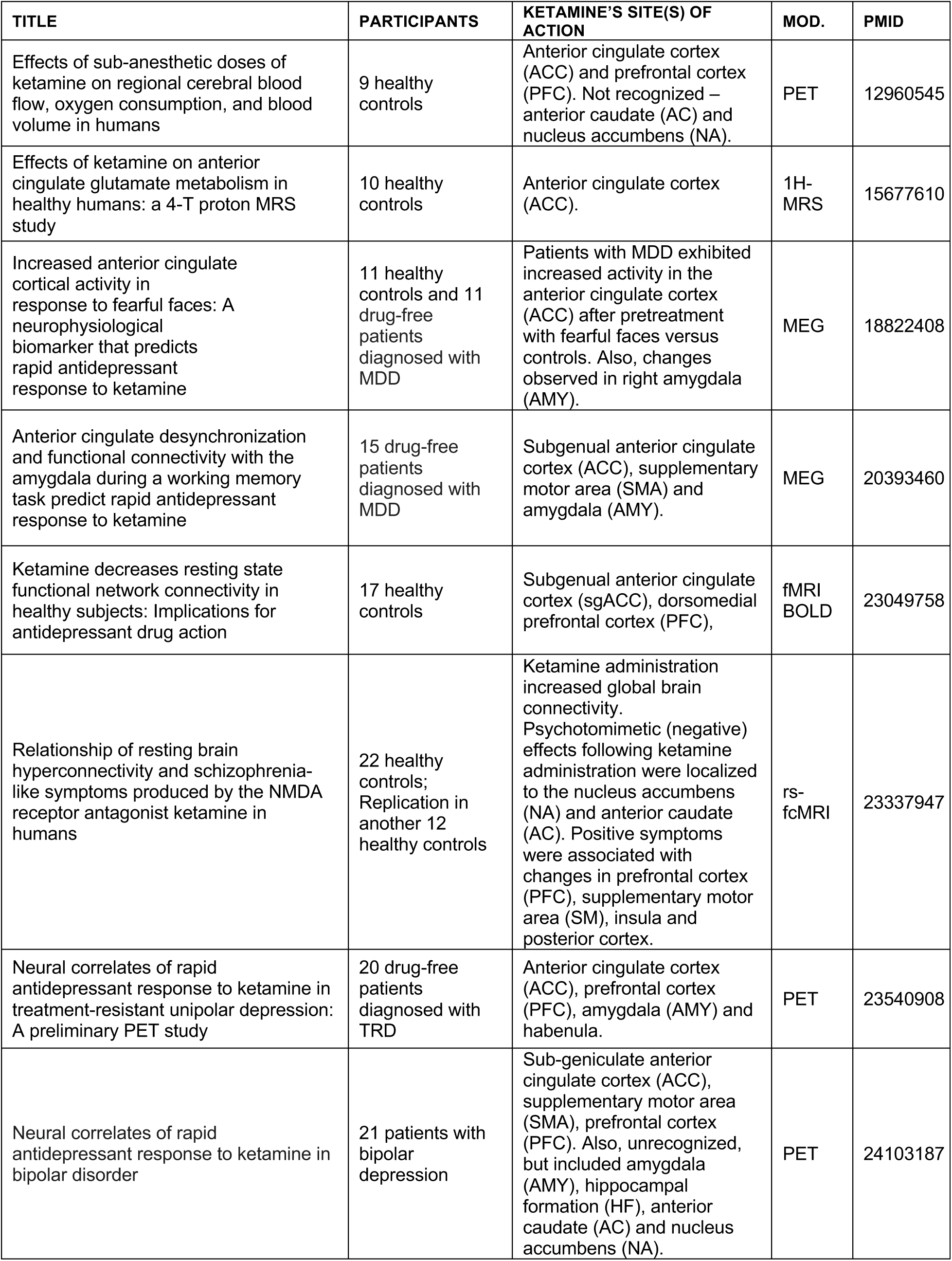
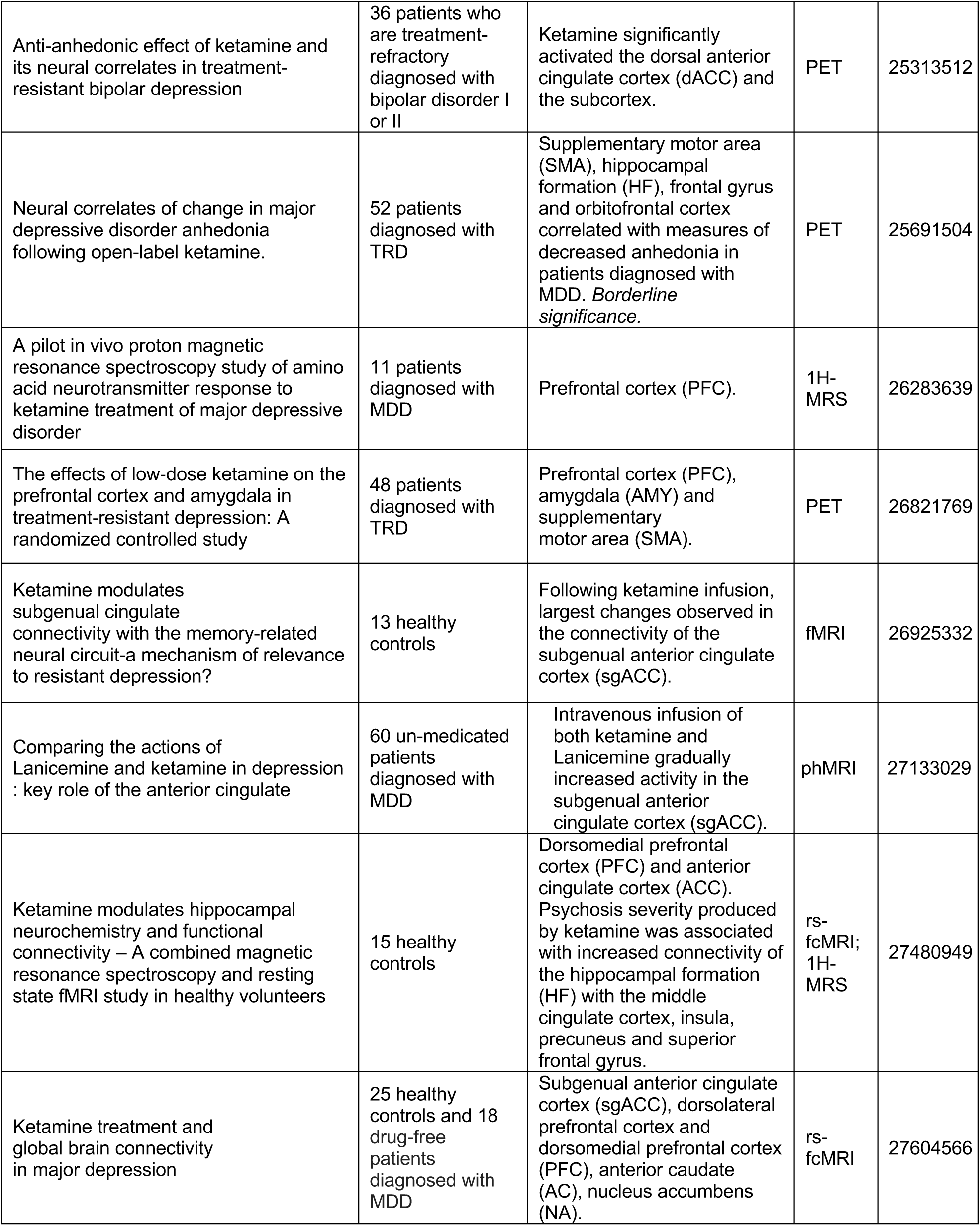
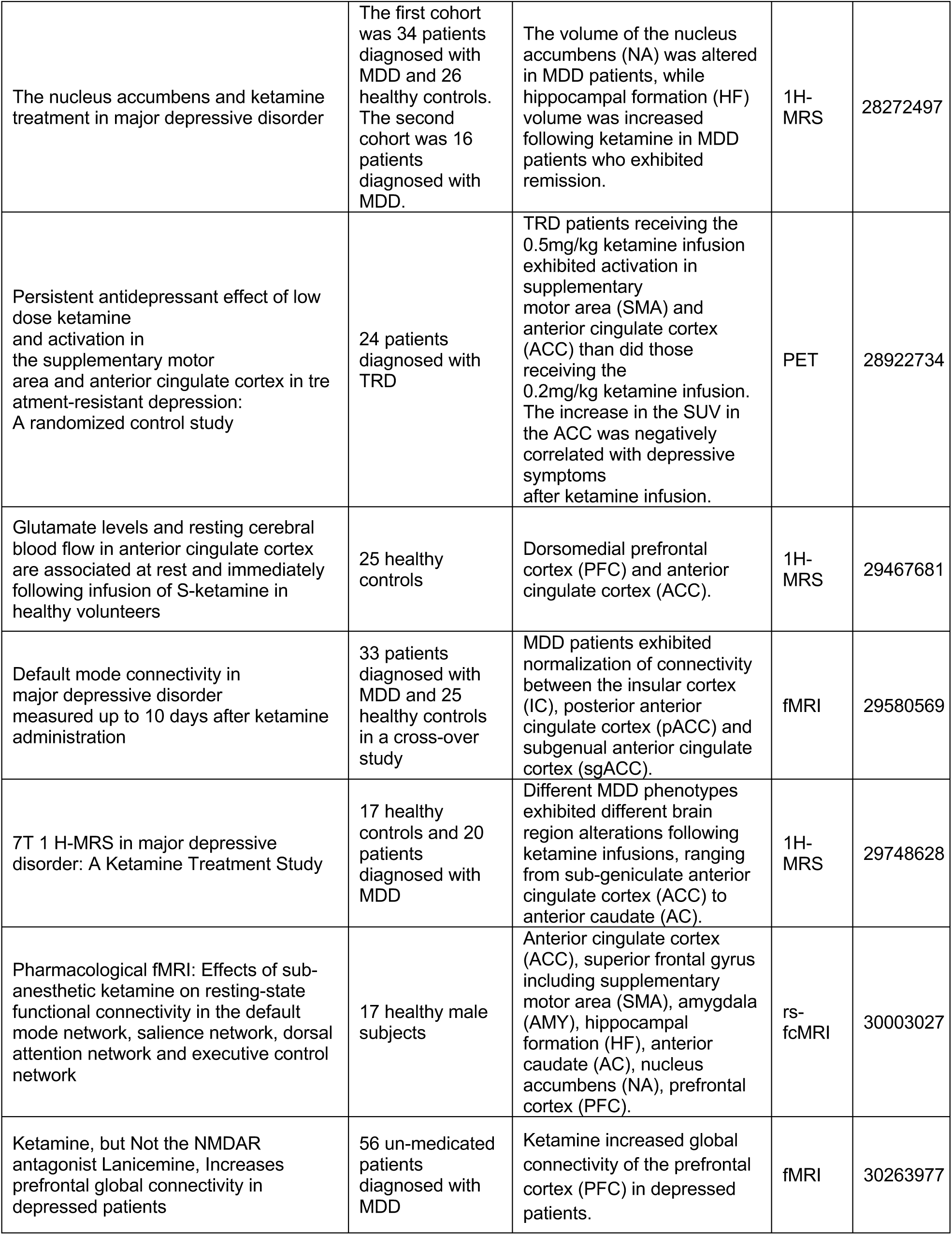
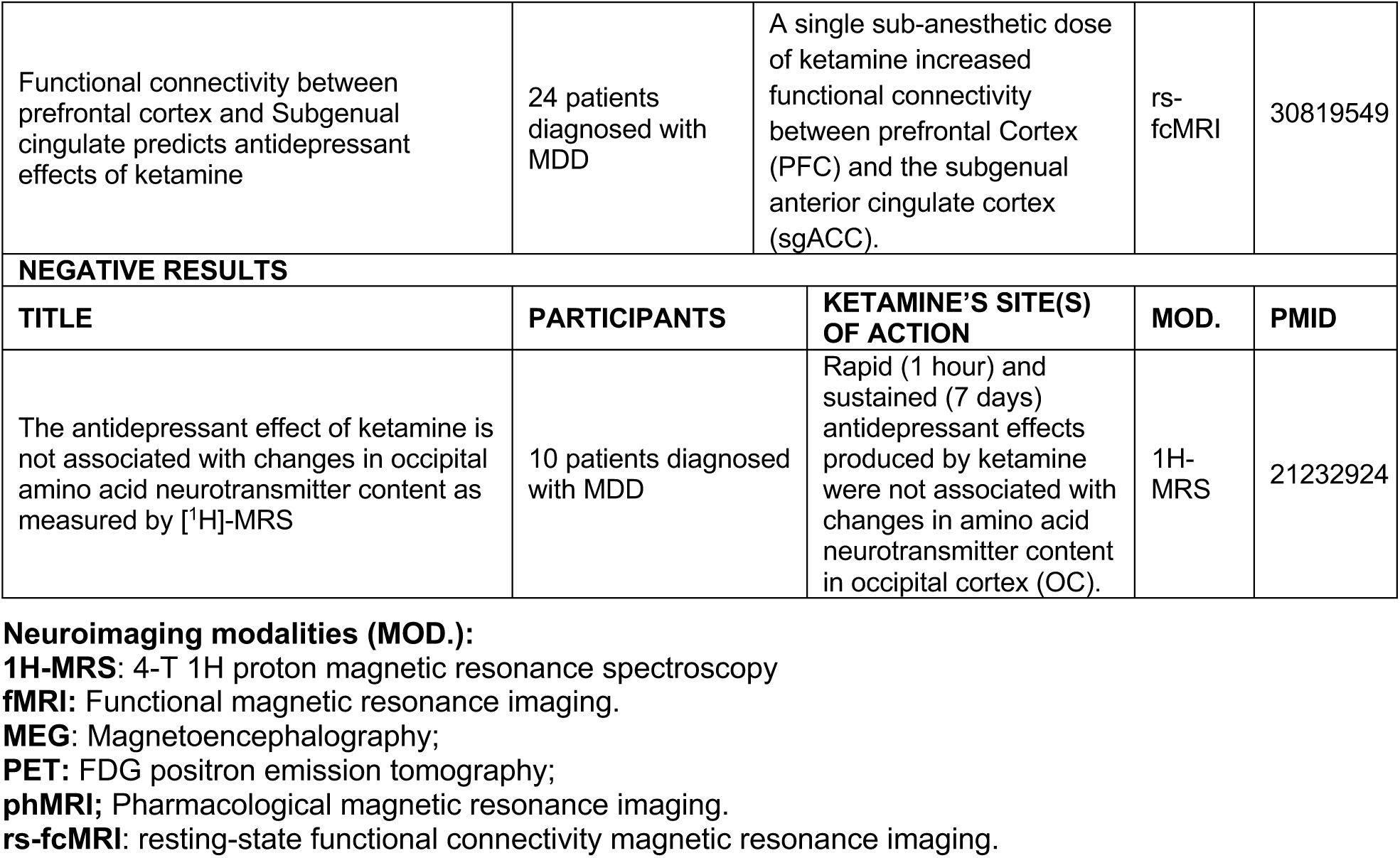
Functional neuroimaging studies used to build consensus map:

**Supplementary Table 2A.** Gene symbols, ketamine glutamate receptor sub-network.

| Gene symbol | Gene name (HGNC) | Location | Protein product |
| --- | --- | --- | --- |
| ACHE | acetylcholinesterase (Cartwright blood group) | Plasma Membrane | enzyme |
| ATF7IP | activating transcription factor 7 interacting protein | Nucleus | transcription regulator |
| ATF7IP2 | activating transcription factor 7 interacting protein 2 | Nucleus | other |
| ATP1A1 | ATPase Na<sup>+</sup>/K<sup>+</sup> transporting subunit alpha 1 | Plasma Membrane | transporter |
| BORCS7 | BLOC-1 related complex subunit 7 | Cytoplasm | other |
| BRD4 | bromodomain containing 4 | Nucleus | kinase |
| CACNA1C | calcium voltage-gated channel subunit alpha1 C | Plasma Membrane | ion channel |
| CACNB1 | calcium voltage-gated channel auxiliary subunit beta 1 | Plasma Membrane | ion channel |
| CACNB2 | calcium voltage-gated channel auxiliary subunit beta 2 | Plasma Membrane | ion channel |
| CACNG2 | calcium voltage-gated channel auxiliary subunit gamma 2 | Plasma Membrane | ion channel |
| CHRM2 | cholinergic receptor muscarinic 2 | Plasma Membrane | G-protein coupled receptor |
| CHRNA3 | cholinergic receptor nicotinic alpha 3 subunit | Plasma Membrane | transmembrane receptor |
| CHRNA5 | cholinergic receptor nicotinic alpha 5 subunit | Plasma Membrane | transmembrane receptor |
| CHRNA7 | cholinergic receptor nicotinic alpha 7 subunit | Plasma Membrane | transmembrane receptor |
| CNR1 | cannabinoid receptor 1 | Plasma Membrane | G-protein coupled receptor |
| DLG3 | discs large MAGUK scaffold protein 3 | Plasma Membrane | kinase |
| DLG4 | discs large MAGUK scaffold protein 4 | Plasma Membrane | kinase |
| DNMT1 | DNA methyltransferase 1 | Nucleus | enzyme |
| EHMT1 | euchromatic histone lysine methyltransferase 1 | Nucleus | transcription regulator |
| GABRA2 | gamma-aminobutyric acid type A receptor alpha2 subunit | Plasma Membrane | ion channel |
| GABRA5 | gamma-aminobutyric acid type A receptor alpha5 subunit | Plasma Membrane | ion channel |
| GAD1 | glutamate decarboxylase 1 | Cytoplasm | enzyme |
| GLRA1 | glycine receptor alpha 1 | Plasma Membrane | ion channel |
| GLRA2 | glycine receptor alpha 2 | Plasma Membrane | ion channel |
| GLRB | glycine receptor beta | Plasma Membrane | ion channel |
| GRIA1 | glutamate ionotropic receptor AMPA type subunit 1 | Plasma Membrane | ion channel |
| GRIA2 | glutamate ionotropic receptor AMPA type subunit 2 | Plasma Membrane | ion channel |
| GRIA4 | glutamate ionotropic receptor AMPA type subunit 4 | Plasma Membrane | ion channel |
| GRIN1 | glutamate ionotropic receptor NMDA type subunit 1 | Plasma Membrane | ion channel |
| GRIN2A | glutamate ionotropic receptor NMDA type subunit 2A | Plasma Membrane | ion channel |
| GRIN2B | glutamate ionotropic receptor NMDA type subunit 2B | Plasma Membrane | ion channel |
| GRIN2C | glutamate ionotropic receptor NMDA type subunit 2C | Plasma Membrane | ion channel |
| GRIN2D | glutamate ionotropic receptor NMDA type subunit 2D | Plasma Membrane | ion channel |
| GRIN3A | glutamate ionotropic receptor NMDA type subunit 3A | Plasma Membrane | ion channel |
| GRIN3B | glutamate ionotropic receptor NMDA type subunit 3B | Plasma Membrane | ion channel |
| HCN1 | hyperpolarization activated cyclic nucleotide gated potassium channel 1 | Plasma Membrane | ion channel |
| HDAC5 | histone deacetylase 5 | Nucleus | transcription regulator |
| MBD1 | methyl-CpG binding domain protein 1 | Nucleus | transcription regulator |
| MPHOSPH8 | M-phase phosphoprotein 8 | Nucleus | transcription regulator |
| NCAM1 | neural cell adhesion molecule 1 | Plasma Membrane | other |
| NOS1 | nitric oxide synthase 1 | Cytoplasm | enzyme |
| NOS2 | nitric oxide synthase 2 | Cytoplasm | enzyme |
| NOS3 | nitric oxide synthase 3 | Cytoplasm | enzyme |
| NQO1 | NAD(P)H quinone dehydrogenase 1 | Cytoplasm | enzyme |
| OPRK1 | opioid receptor kappa 1 | Plasma Membrane | G-protein coupled receptor |
| OPRM1 | opioid receptor mu 1 | Plasma Membrane | G-protein coupled receptor |
| ROBO2 | roundabout guidance receptor 2 | Plasma Membrane | transmembrane receptor |
| SETDB1 | SET domain bifurcated histone lysine methyltransferase 1 | Nucleus | enzyme |
| SHANK2 | SH3 and multiple ankyrin repeat domains 2 | Plasma Membrane | other |
| SIGMAR1 | sigma non-opioid intracellular receptor 1 | Plasma Membrane | transmembrane receptor |
| SLC6A9 | solute carrier family 6 member 9 | Plasma Membrane | transporter |
| TASOR | transcription activation suppressor | Nucleus | other |
| TOGARAM2 | TOG array regulator of axonemal microtubules 2 | Other | other |
| TRIM28 | tripartite motif containing 28 | Nucleus | transcription regulator |
| ZNF274 | zinc finger protein 274 | Nucleus | transcription regulator |

**Supplementary Table 2B.** Gene symbols, ketamine neuroplasticity sub-network

| Gene symbol | Gene name (HGNC) | Location | Protein product |
| --- | --- | --- | --- |
| ARC | activity regulated cytoskeleton associated protein | Cytoplasm | other |
| ASCL1 | achaete-scute family bHLH transcription factor 1 | Nucleus | transcription regulator |
| BDNF | brain derived neurotrophic factor | Extracellular Space | growth factor |
| BDNF-AS | BDNF antisense RNA | Other | other |
| CAMK2A | calcium/calmodulin dependent protein kinase II alpha | Cytoplasm | kinase |
| CDKN1A | cyclin dependent kinase inhibitor 1A | Nucleus | kinase |
| CREM | cAMP responsive element modulator | Nucleus | transcription regulator |
| CUX2 | cut like homeobox 2 | Nucleus | transcription regulator |
| DCC | DCC netrin 1 receptor | Plasma Membrane | transmembrane receptor |
| DRD2 | dopamine receptor D2 | Plasma Membrane | G-protein coupled receptor |
| EEF2K | eukaryotic elongation factor 2 kinase | Cytoplasm | kinase |
| FMR1 | fragile X mental retardation 1 | Cytoplasm | translation regulator |
| GDAP1L1 | ganglioside induced differentiation associated protein 1 like 1 | Other | other |
| GRM5 | glutamate metabotropic receptor 5 | Plasma Membrane | G-protein coupled receptor |
| HOMER1 | homer scaffold protein 1 | Plasma Membrane | other |
| HTR1B | 5-hydroxytryptamine receptor 1B | Plasma Membrane | G-protein coupled receptor |
| HTR2A | 5-hydroxytryptamine receptor 2A | Plasma Membrane | G-protein coupled receptor |
| KLF6 | Kruppel like factor 6 | Nucleus | transcription regulator |
| LIN7C | lin-7 homolog C, crumbs cell polarity complex component | Cytoplasm | other |
| ENSG00000251574 |  | Nucleus | RNA gene |
| MEF2D | myocyte enhancer factor 2D | Nucleus | transcription regulator |
| MYO6 | myosin VI | Cytoplasm | other |
| MYT1L | myelin transcription factor 1 like | Nucleus | transcription regulator |
| NEUROD1 | neuronal differentiation 1 | Nucleus | transcription regulator |
| NEUROD2 | neuronal differentiation 2 | Nucleus | transcription regulator |
| NHLH2 | nescent helix-loop-helix 2 | Nucleus | transcription regulator |
| NMB | neuromedin B | Extracellular Space | other |
| NSMF | NMDA receptor synaptonuclear signaling and neuronal migration factor | Extracellular Space | other |
| NTRK2 | neurotrophic receptor tyrosine kinase 2 | Plasma Membrane | kinase |
| PTEN | phosphatase and tensin homolog | Cytoplasm | phosphatase |
| PTGS2 | prostaglandin-endoperoxide synthase 2 | Cytoplasm | enzyme |
| RAC1 | Rac family small GTPase 1 | Plasma Membrane | enzyme |
| RASGRF2 | Ras protein specific guanine nucleotide releasing factor 2 | Cytoplasm | other |
| RHOA | ras homolog family member A | Cytoplasm | enzyme |
| ROBO2 | roundabout guidance receptor 2 | Plasma Membrane | transmembrane receptor |
| ENSG00000225960 |  | Nucleus | RNA gene |
| SEMA3A | semaphorin 3A | Extracellular Space | other |
| SHANK1 | SH3 and multiple ankyrin repeat domains 1 | Cytoplasm | other |
| SHANK2 | SH3 and multiple ankyrin repeat domains 2 | Plasma Membrane | other |
| SHANK3 | SH3 and multiple ankyrin repeat domains 3 | Plasma Membrane | other |
| SLC22A15 | solute carrier family 22 member 15 | Other | transporter |
| SLC6A2 | solute carrier family 6 member 2 | Plasma Membrane | transporter |
| SLIT1 | slit guidance ligand 1 | Extracellular Space | other |
| SLIT2 | slit guidance ligand 2 | Extracellular Space | other |
| SNAP25 | synaptosome associated protein 25 | Plasma Membrane | transporter |
| SYN1 | synapsin I | Plasma Membrane | transporter |
| SYN2 | synapsin II | Plasma Membrane | other |
| SYT3 | synaptotagmin 3 | Cytoplasm | transporter |
| TBR1 | T-box, brain 1 | Nucleus | transcription regulator |
| TCF4 | transcription factor 4 | Nucleus | transcription regulator |

**Supplementary Table 2C.** C Gene symbols, ketamine pharmacokinetic receptor sub-network

| Gene symbol | Gene name (HGNC) | Location | Protein product |
| --- | --- | --- | --- |
| ANAPC2 | anaphase promoting complex subunit 2 | Nucleus | other |
| CYP2A6 | cytochrome P450 family 2 subfamily A member 6 | Cytoplasm | enzyme |
| CYP2B6 | cytochrome P450 family 2 subfamily B member 6 | Cytoplasm | enzyme |
| CYP3A4 | cytochrome P450 family 3 subfamily A member 4 | Cytoplasm | enzyme |
| ESR1 | estrogen receptor 1 | Nucleus | ligand-dependent nuclear receptor |
| TCERG1 | transcription elongation regulator 1 | Nucleus | transcription regulator |

**Supplementary Table 3.** **Ketamine binding affinity studies**. Affinity of ketamine to various CNS targets as measured by displacement of radiolabeled ligands or radioligand assays. **Molecules in bold text were added to the ketamine network based on the high affinity binding of ketamine**. CHO: Chinese hamster ovary cells; PubMed ID numbers refers to where the original studies can be found at the PubMed web site, National Library of Medicine **(65)**.

| TARGET | SPECIES | BINDING AFFINITY / INHIBITION CONSTANT | STEREOSELECTIVITY | PMID |
| --- | --- | --- | --- | --- |
| **AMPA:**<br>**GRIA1,**<br>**GRIA2,**<br>**GRIA4** | Human | $K_i = 7 \pm 0.5 \mu\text{M}$ (SK), $24.5 \pm 2.0 \mu\text{M}$ (RK) | SK has 3.5-fold affinity of RK | 29945898 |
| **NMDAR:**<br>**GRIN1,**<br>**GRIN2A,**<br>**GRIN2B,**<br>**GRIN2C,**<br>**GRIN2D,**<br>**GRIN3A,**<br>**GRIN3B** | Human | $K_i = 0.7 \pm 0.3 \mu\text{M}$ (SK), $2.3 \pm 0.3 \mu\text{M}$ (RK) | SK has 3-fold affinity of RK | 29945898 |
| **NMDAR:**<br>**GRIN1,**<br>**GRIN2A,**<br>**GRIN2B,**<br>**GRIN2C,**<br>**GRIN2D,**<br>**GRIN3A,**<br>**GRIN3B** | Human | $K_i = 0.06 \pm 0.1 \mu\text{M}$ | Not measured | 1311263 |
| **CACNA1C** | Rat | $IC_{50} = 9.2 \mu\text{M}$ | Not measured | 29945898 |
| **CHRM2** | Human | $K_i = 7.4 \pm 4.5 \mu\text{M}$ | No observed difference | 29945898 |
| **CNR1** | Human | $K_i = 42 \pm 5 \mu\text{M}$ | Not measured | 24093505 |
| **CHRNA3** | Human | $IC_{50} = 50 \mu\text{M}$ | Not measured | 29945898 |
| **CHRNA5** | Human | $IC_{50} = 5 \mu\text{M}$ | Not measured | 29945898 |
| **CHRNA7** | Human | $K_i = 12 \pm 0.8 \mu\text{M}$ | No observed difference | 29945898 |
| **DRD2** | Human | $K_i = 12 \pm 0.8 \mu\text{M}$ | Not measured | 12232776 |
| **DRD2** | Rat | $K_i = 1 \pm 0.2 \mu\text{M}$ | Not measured | 30034974 |
| **ESR1** | Human | $K_i = 94 \pm 8 \mu\text{M}$ | No observed difference | 29945898 |
| GABRA1 | Human | $K_i = 134 \pm 29 \mu\text{M}$ | Not measured | 29945898 |
| GABBR1 | Human | $K_i = 144 \pm 32 \mu\text{M}$ | Not measured | 29945898 |
| **GABRA2** | Guinea pig | $K_i = 1 \pm 0.7 \mu\text{M}$ (RK), $12.2 \pm 1.2 \mu\text{M}$ (SK) | RK has 12-fold affinity of SK | 29945898 |
| **GABRA5** | Rat | $K_i = 10 \pm 1.1 \mu\text{M}$ | No observed difference | 29945898 |
| **GLRB** | Human | $K_i = 27 \pm 7.2 \mu\text{M}$ | Not measured | 29945898 |
| **GRM5** | Rat | $K_i = 7 \pm 0.62 \mu\text{M}$ | Not measured | 30034974 |
| **HCN1** | Rat PC12 cells | $K_i = 0.035 \pm 0.3 \mu\text{M}$ (RK), $54 \pm 3.7 \mu\text{M}$ (SK) | RK has 5000-fold affinity of SK | 29945898 |
| **HCN1** | Human | $EC_{50} = 4.1 \mu\text{M} - 7.1 \mu\text{M}$ (SK) | Not measured | 29945898 |
| HTR3A | Human | $IC_{50} = 910 \pm 30 \mu\text{M}$ | Not measured | 10754635 |
| **HTR2A** | Human | $K_i = 19 \mu\text{M}$ (RK), $135 \mu\text{M}$ (SK), | RK has 7-fold affinity of SK | 29945898 |
| **HTR1A** | Human | $IC_{50} = 46 \pm 1.4 \mu\text{M}$ | Not measured | 29945898 |
| **HTR1B** | Human | $K_i = 6 \pm 0.2 \mu\text{M}$ | Not measured | 29945898 |
| **OPRM1** | Guinea pig | $K_i = 11 \mu\text{M}$ (SK), $28 \mu\text{M}$ (RK) | SK has 2.5-fold affinity of RK | 8835358 |
| OPRD1 | Guinea pig | $K_i = 130 \mu\text{M}$ (SK), $130 \mu\text{M}$ (RK) | No significant difference | 8835358 |
| **OPRK1** | CHO | $K_i = 4.2 \pm 0.03 \mu\text{M}$ (SK), $4.6 \pm 0.04 \mu\text{M}$ (RK) | No significant difference | 9915326 |
| **SIGMAR1** | Rat brain | $K_i = 19 \mu\text{M}$ (RK), $200 \mu\text{M}$ (SK) | RK has 10-fold affinity of SK | 29945898 |
| **SLC6A2** | Human | $K_i = 67 \pm 26 \mu\text{M}$ | Not measured | 9523822 |
| **SLC6A3** | Human | $K_i = 46.9 \mu\text{M}$ (SK), $390 \mu\text{M}$ (RK) | SK has 8-fold affinity of RK | 10553955 |
| **SLC6A4** | Rat | $K_i = 162 \pm 28 \mu\text{M}$ | Not measured | 9523822 |

**Supplementary Fig. 1.**
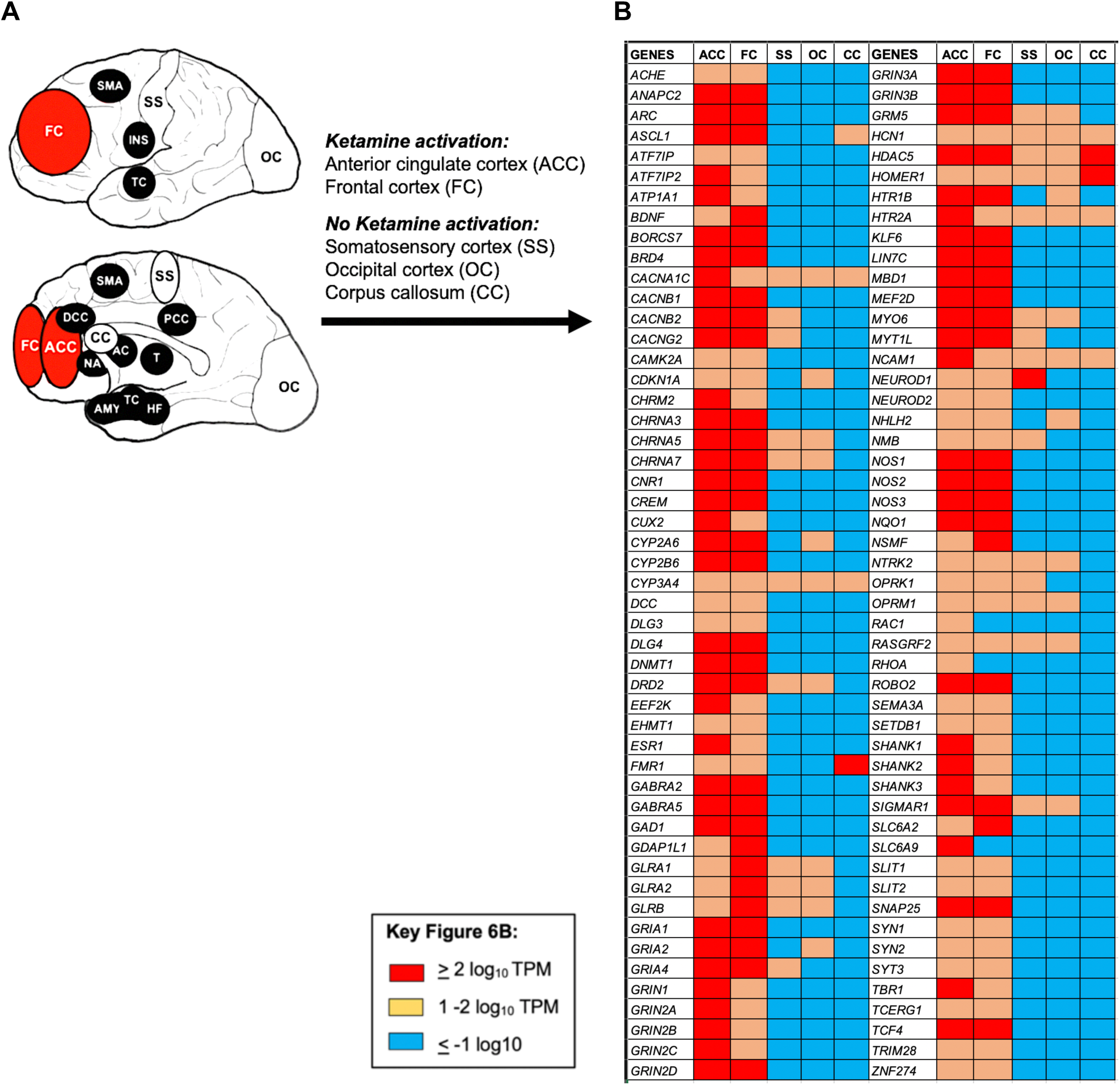
Consensus between ketamine pharmacogenomic pathway gene expression in human brain and regions where ketamine exerts a rapid antidepressant effect derived from functional imaging studies (Supplementary Table 1). **(A)** Human brain regions impacted by ketamine projected onto lateral (top) and medial (bottom) surfaces of the brain. Black indicates ketamine activation regions and dark red indicates those CNS areas first activated by the drug. **(B)** Expression of 100 out of 107 genes in the ketamine pharmacogenomic pathway whose expression could be determined. Abbreviations: **ACC**: Anterior cingulate cortex; **AC**: Anterior caudate; **AMY**: Amygdala; **CC**: Corpus callosum; **dACC**: Dorsal anterior cingulate cortex; **FC**: Frontal cortex; **HF**: Hippocampal formation; **INS**: Insular cortex; **NA**: Nucleus accumbens; **OC**: Occipital cortex; **pCC**: Posterior cingulate cortex; **PFC**: Prefrontal cortex; **SMA**: Supplementary motor area; **Supplementary Figure 3**. Examples of *in situ* hybridization of selected super-pathway genes that are ketamine PD targets, from the human brain atlas of the Allen Brain Science Institute **(62) sgACC**: Subgenual anterior cingulate cortex; **SS**: Somatosensory cortex; **T**: Thalamus; **TC**: Temporal cortex; **TPM**: Transcripts per million.

**Supplementary Figure 2.**
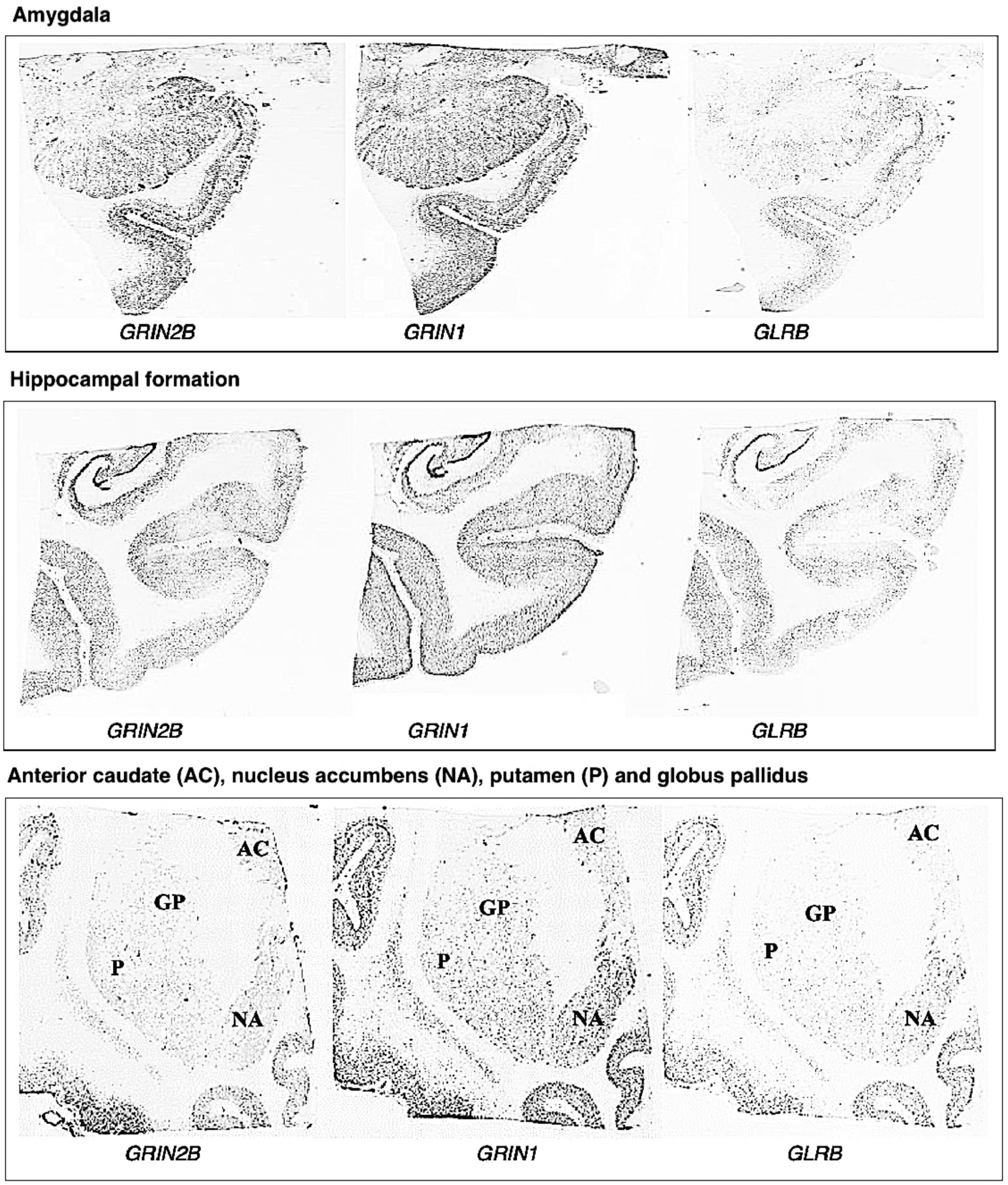
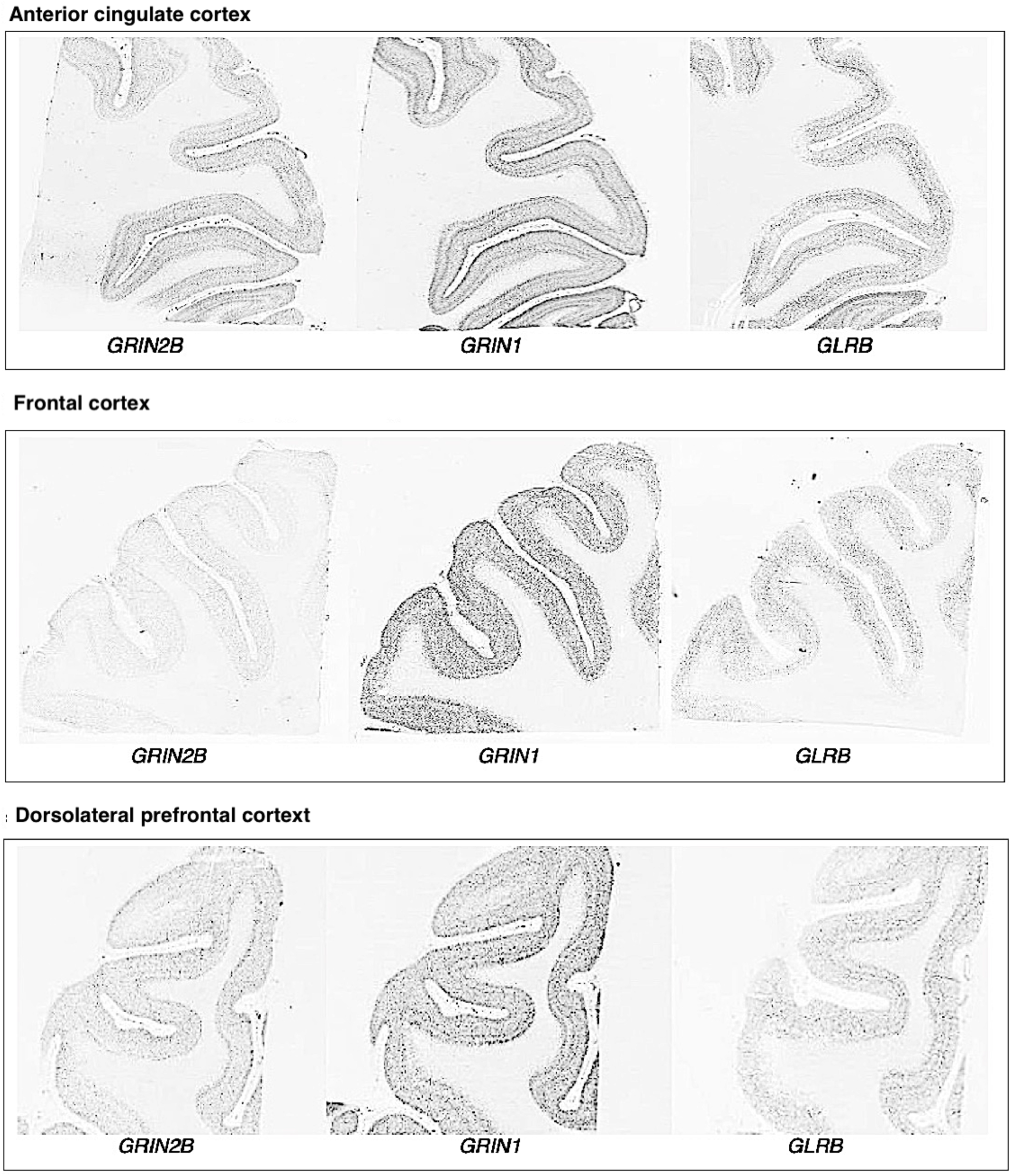
Examples of *in situ* hybridization of selected super-pathway genes that are ketamine PD targets, from (62)

**Additional Figure captions:**

**Supplementary Figure 4**. GWAS SNPs from the ketamine glutamate receptor sub-network

**Supplementary Figure 5**. GWAS SNPs from the ketamine neuroplasticity sub-network

